# Conformational Dynamics and Catalytic Backups in a Hyper-Thermostable Engineered Archaeal Protein Tyrosine Phosphatase

**DOI:** 10.1101/2025.03.26.645524

**Authors:** Dariia Yehorova, Nikolas Alansson, Ruidan Shen, Joshua M. Denson, Michael Robinson, Valeria A. Risso, Nuria Ramirez Molina, J. Patrick Loria, Eric A. Gaucher, Jose M. Sanchez-Ruiz, Alvan C. Hengge, Sean J. Johnson, Shina C. L. Kamerlin

## Abstract

Protein tyrosine phosphatases (PTPs) are a family of enzymes that play important roles in regulating cellular signaling pathways. The activity of these enzymes is regulated by the motion of a catalytic loop that places a critical conserved aspartic acid side chain into the active site for acid-base catalysis upon loop closure. These enzymes also have a conserved phosphate binding loop that is typically highly rigid and forms a well-defined anion binding nest. The intimate links between loop dynamics and chemistry in these enzymes make PTPs an excellent model system for understanding the role of loop dynamics in protein function and evolution. In this context, archaeal PTPs, which have evolved in extremophilic organisms, are highly understudied, despite their unusual biophysical properties. We present here an engineered chimeric PTP (ShufPTP) generated by shuffling the amino acid sequence of five extant hyperthermophilic archaeal PTPs. Despite ShufPTP’s high sequence similarity to its natural counterparts, ShufPTP presents a suite of unique properties, including high flexibility of the phosphate binding P-loop, facile oxidation of the active site cysteine, mechanistic promiscuity, and most notably, hyperthermostability, with a denaturation temperature likely >130 °C (>8 °C higher than the highest recorded growth temperature of any archaeal strain). Our combined structural, biochemical, biophysical and computational analysis provides insight both into how small steps in evolutionary space can radically modulate the biophysical properties of an enzyme, and showcase the tremendous potential of archaeal enzymes for biotechnology, to generate novel enzymes capable of operating under extreme conditions.

**Table of Contents Graphic:** 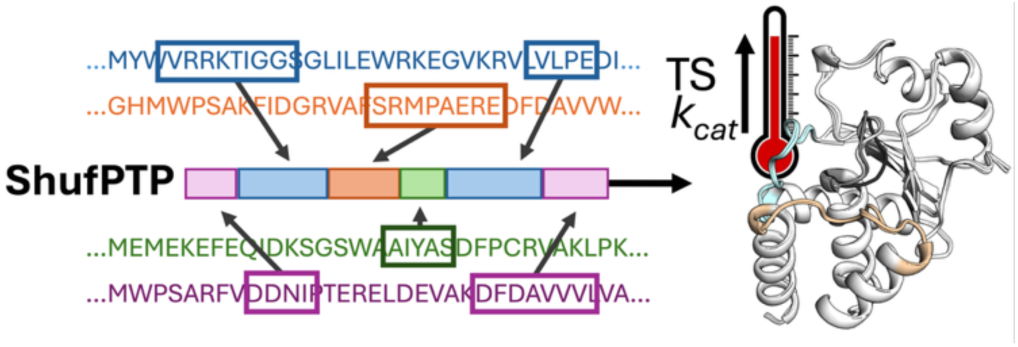

## Introduction

Enzymes are exquisite catalysts,^1^ and there remains tremendous interest in exploiting tailored enzymes for use in biotechnology and beyond.^2-5^ In this context, there is increasing evidence that conformational dynamics plays an important role in the emergence of new enzyme activities, and in the evolutionary optimization of enzyme selectivity and activity.^6-34^ In particular, enzyme active site loops are highly flexible, and evidence shows that evolutionary conformational modulation of their dynamical behavior can translate into control of enzyme specificity, activity, and even the pH dependency of catalysis.^35^ Understanding how active site loop dynamics is evolutionarily and allosterically regulated, and how it is linked to the chemical step of catalysis, is thus important for our understanding of the factors shaping new enzyme functions and for protein engineering to control protein activity, selectivity, and biophysical properties.^35, 36^

Protein tyrosine phosphatases are an excellent model system for probing the links between loop dynamics and the evolution of enzyme activity. These genetically diverse enzymes^37^ share a common core structure, chemical mechanism, and enzymatic transition states.^38^ Yet, their catalytic rates vary by orders of magnitude,^39^ reflecting the variety of regulatory roles they play *in vivo*. Structurally, PTPs share a number of catalytic loops that decorate their active sites (**Figure 1**): (1) the highly conserved phosphate-binding P-loop; (2) a highly mobile acid loop (the WPD/IPD loop), which carries a critical aspartic acid and undergoes a substantial conformational change (∼10Å) between catalytically inactive “open” and active “closed” conformations; as well as (3) additional Q- and E-loops, which carry catalytically and structurally important residues.^31^ From a dynamical perspective, it is noteworthy that, unlike many enzymes that are regulated by catalytic loop motion, the acid loop of PTPs does not form a lid over the active site. Rather, it plays a key chemical role, positioning the conserved Asp side chain on the loop in an optimal position for catalysis (**Figure 1**). Thus, unlike say TIM-barrel proteins (which are frequently decorated by mobile catalytic loops^40^), the PTP acid loop does not form an active site cage, but rather the role of loop motion is primarily chemical, and substrate/product can diffuse in and out of the active site from an acid loop closed position.^41^

**Figure 1.**
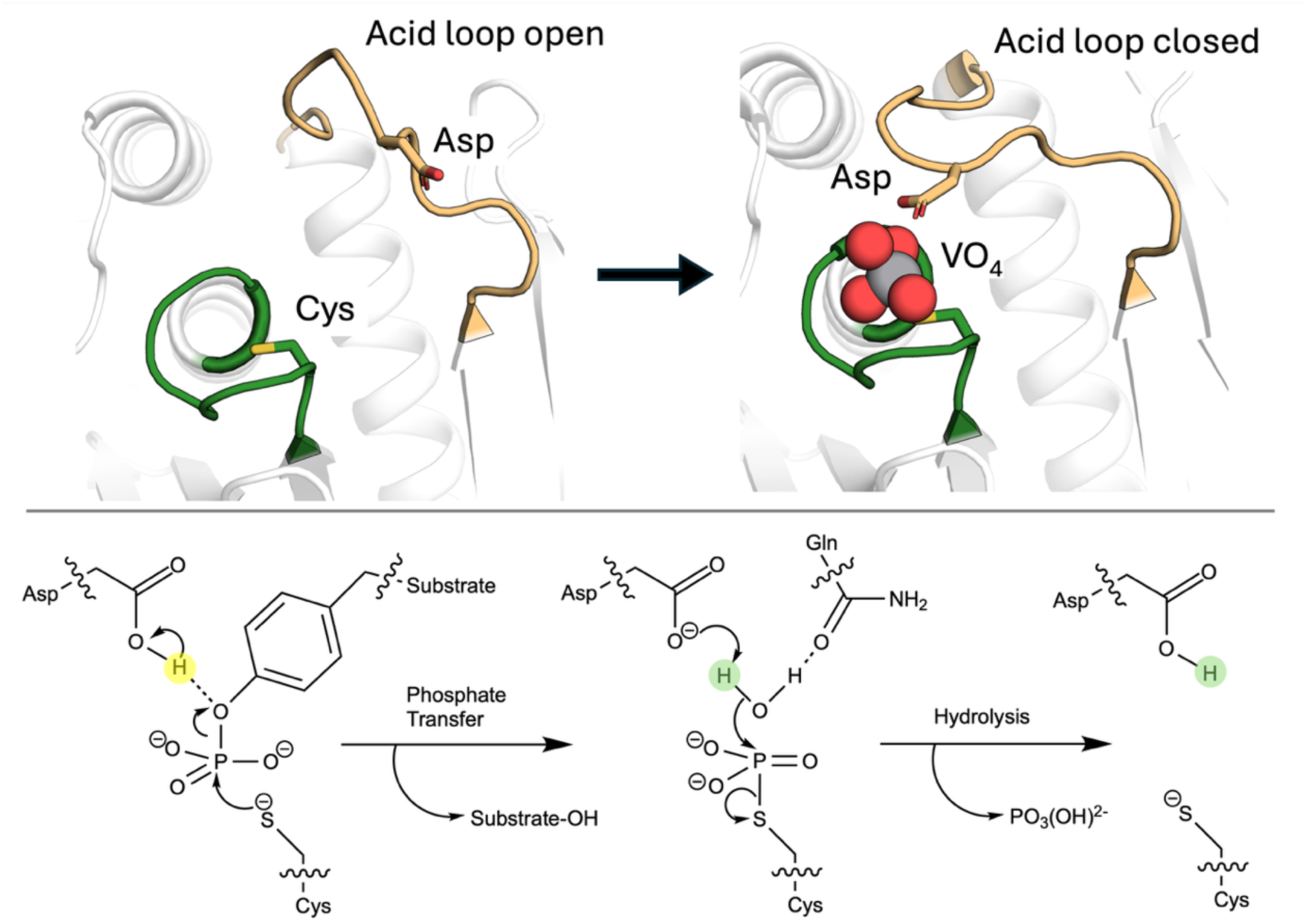
Overview of the PTP catalytic mechanism. (**Top**) Acid loop open and closed conformations of YopH, in its unliganded form and in complex with a VO_4_ transition state analog (PDB IDs: 1YPT^42^ and 2I42,^43^ respectively). The position of the catalytic aspartic acid is highlighted. (**Bottom**) Two-step mechanism catalyzed by PTPs. Adapted from Ref. ^44^. Available under a CC-BY license. Copyright © 2024 The Authors.

In the best characterized PTPs, PTP1B^45^ and YopH,^46^ burst kinetics indicate that hydrolysis is rate-limiting. However, NMR studies of these PTPs have shown correlation between the rates of loop motion and phosphotyrosine cleavage kinetics.^47^ This has been supported by computational, biochemical and structural characterization, suggesting a direct link between loop dynamics, turnover rates, and pH-dependency, in wild-type PTP1B and YopH and variants.^31-33, 39, 47-49^ Furthermore, computational studies indicate that the acid loop of PTPs is conformationally plastic, sampling a diversity of open conformations.^31, 33^ This is an important observation, as evidence suggests that increased activity among the PTP superfamily is directly linked to both destabilization of the acid loop open conformation, and stabilization of the catalytically active loop-closed conformation.^50^

Curiously, while the phosphate-binding P-loop is relatively rigid and structurally conserved in many enzymes,^31, 51-55^ archaeal PTPs are a notable exception, with evidence for temperature-dependent transitions between active and inactive conformations (**Figure 2**).^55, 56^ Evolutionary studies suggest that the phosphorylation/dephosphorylation machinery of eukaryotes and prokaryotes evolved from archaea,^57, 58^ whose PTPs are less characterized than their eukaryotic and prokaryotic counterparts. Therefore, better understanding of the unusual dynamical properties of catalytic loops in archaeal PTPs will aid in understanding the links between loop dynamics and catalysis in the PTP superfamily more broadly, and in enzymes regulated by catalytic loop motion more generally.^36^

**Figure 2.**
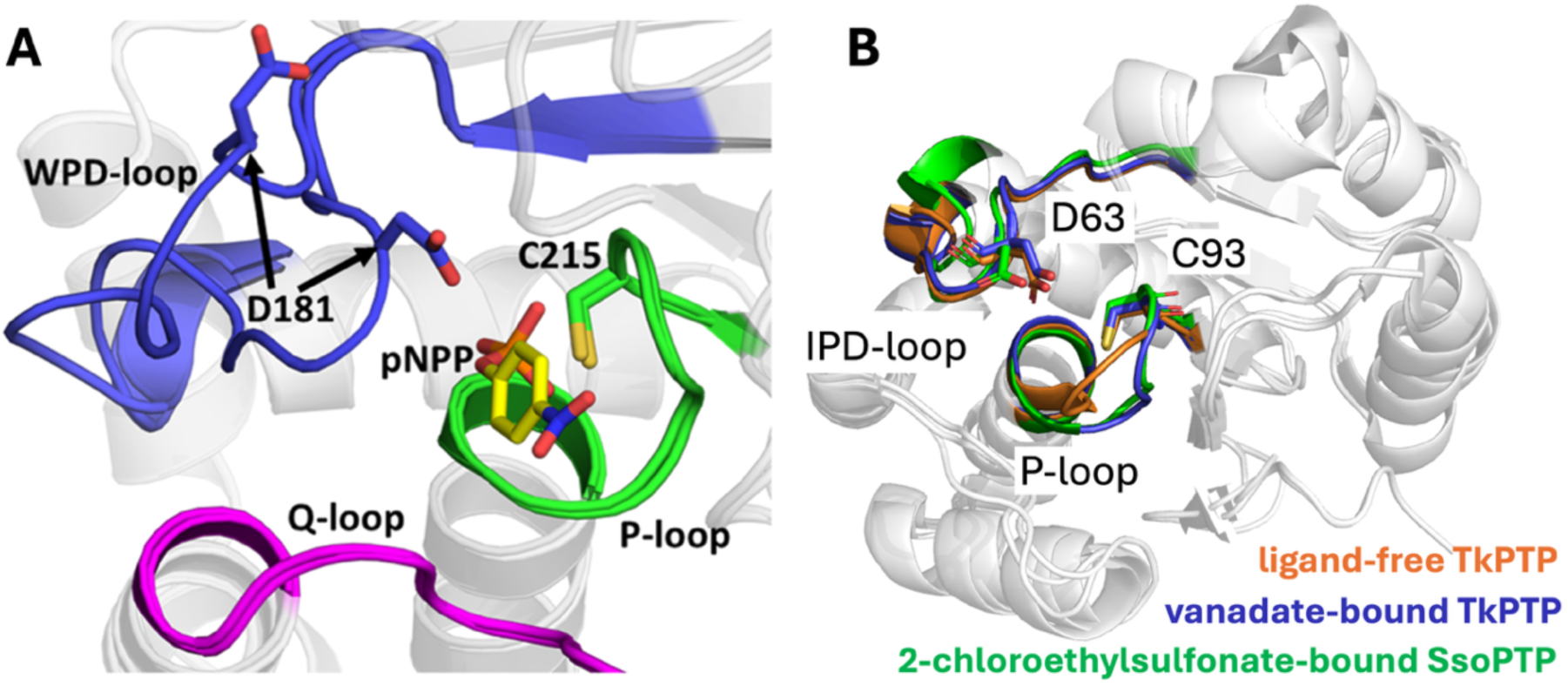
Key loops decorating PTP active sites. (**A**) A comparison of the open and closed conformations of the acid loop (WPD-loop) of the archetypal member of the protein tyrosine phosphatase superfamily, PTP1B, highlighting also the Q- and P-loops. (**B**) In archaeal PTPs, the WPD-loop is replaced by an IPD-loop. Shown here is an overlay of the IPD- and P-loops of the archaeal PTPs, SsoPTP (PDB ID: 7MPD^55^) and TkPTP (PDB IDs: 5Z5A and 5Z59^56^), in both their unliganded and liganded forms. In both archaeal PTPs, the acid loop is closed and in the same position in all structures, but the phosphate binding P-loop is flexible and takes on multiple conformations (note that only one conformation of SsoPTP’s P-loop could be captured structurally, the existence of the other was determined through NMR spectroscopy^55^). Panel A was originally published in ref. ^31^ under a CC-BY license. Published by the American Chemical Society. Copyright © 2021 The Authors.

DNA shuffling is an evolutionary protein engineering technique. It can be used to expand protein functional diversity, manipulate protein biophysical properties, and has a well-established history as a tool for protein engineering.^56, 59-61^ Although best known in experimental synthetic biology, random sequence shuffling is an important tool in computational biology in order to evaluate the statistical significance of a given biological sequence^62^ (for relevant examples, see *e.g*., refs. ^62-68^). Here, we have exploited this technique as an engineering tool by computationally shuffling the amino acid sequences of five thermophilic archaeal PTP sequences to generate a synthetic chimeric archaeal PTP, ShufPTP, with highly unusual biochemical and biophysical properties. ShufPTP shares 90% sequence similarity with its closest extant counterpart, a PTP from the thermophilic anaerobic archaeon, *Thermococcus gorgonarius* (Genbank ID: WP_088884773.1, TgPTP), from a family of archaea typically found in hydrothermal vents.^69^ Computational, structural, biochemical and biophysical characterization shows that ShufPTP is (1) hyperthermostable (*T*_M_ > 130 °C), (2) has a mobile phosphate-binding P-loop that takes on multiple configurations, (3) is mechanistically promiscuous, exploiting a backup catalytic mechanism, and (4) has a nucleophilic cysteine that is readily amenable to oxidation, a property that has been suggested to be important for the regulation of PTP activity.^70-76^ Glimmers of each of these properties are seen in extant archaeal PTPs such as TkPTP^56^ and SsoPTP,^55^ but not to the extent observed in the shuffled protein. This has important implications for protein engineering, suggesting an untapped pool of enzyme scaffolds among archaeal enzymes from which to develop novel enzymes having extreme properties (in particular thermostability) for biocatalytic purposes.

## Results and Discussion

### Random Sequence Shuffling of Thermophilic Archaeal PTPs

ShufPTP was obtained by aligning the 5 extant PTP sequences shown in **Figure S1,** based on MUltiple Sequence Comparison by Log-Expectation (MUSCLE)^77^ of the progenitor sequences, using the EMBL-EBI job dispatcher.^78^ For each position, analyzed, when a site is conserved by a single amino acid, that residue is selected for the random sequence. When a site contains multiple amino acid residues, a single amino acid is selected for the random sequence. The selection of the residue is unweighted. This is equivalent to randomly selecting a residue from the posterior probability distribution to sample sequence space (as previously shown in ref. ^79^).

These specific sequences were selected based on their initial annotation as PTPs, followed by manual validation comparing the sequences and their predicted structures (based on structure prediction using the AlphaFold server^80^) to previously characterized PTPs (TkPTP,^56, 81^ SsoPTP,^55^ and VHZ^82, 83^). They were also chosen because of the high growth temperatures observed for the selected organisms, which range, depending on organism, between 50-95°C,^84-88^ and with optimal temperatures in the range of 80 - 88 °C for *Thermococcus gorgonarius*,^84^ 80 °C for *Thermococcus celericrescens*,^87^ 85 °C for *Thermococcus siculi*,^85^ 85 °C for *Thermococcus cleftensis*,^88^ and 88 °C for *Thermococcus radiotolerans*.^86^ Enzyme thermal stability is usually (but not always) correlated with the corresponding growth temperatures,^89^ and can even be used as meaningful training data for the deep learning algorithms that predict thermostability.^90^ The high growth temperatures of the parent organisms for the PTPs shown in **Figure S1** and used for the amino acid sequence shuffling thus in turn imply likely high thermal stability for the corresponding PTPs. For further comparison, **Figure 3** shows sequence alignment between ShufPTP, three archaeal PTPs (TgPTP, TkPTP and SsoPTP, the first two of which were used for sequence shuffling), and the human and bacterial PTPs PTP1B and YopH, which are two of the best-studied PTPs.^41, 91^.

**Figure 3.**
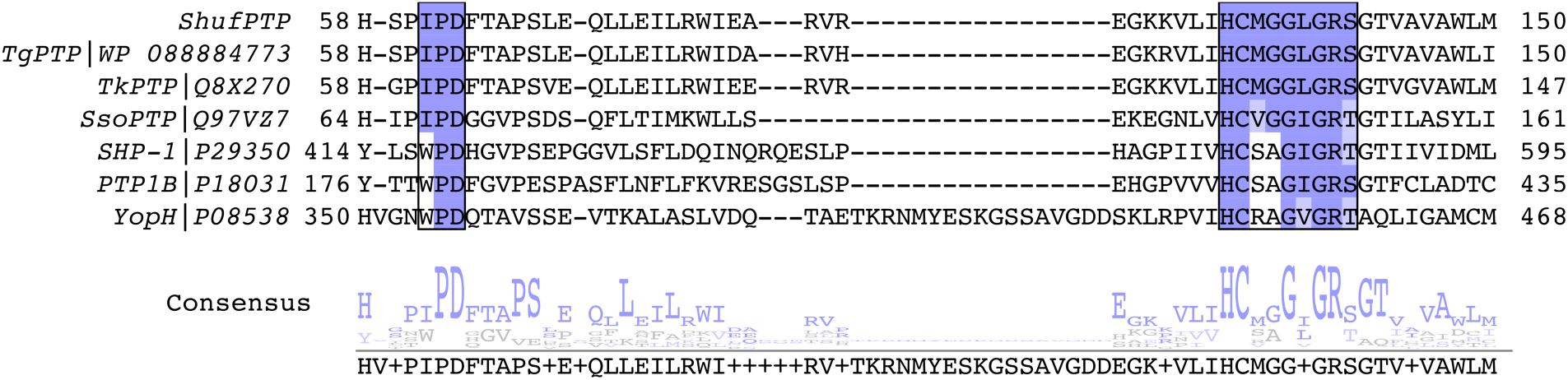
Sequence conservation demonstrates the archaeal nature of ShufPTP. Multiple sequence alignment of characterized PTPs with the synthetic PTP ShufPTP highlights its similarities to thermophilic archaeal PTPs. Shown here is a portion of the sequence alignment among two archaeal PTPs from *Thermococci*, TgPTP and TkPTP,^56^ a third archaeal PTP SsoPTP,^55^ the human PTPs SHP-1 and PTP1B, and the bacterial PTP YopH, as well as the corresponding consensus sequence. Sequence alignment was performed using T-Coffee.^95^ The conserved IPD- or WPD-regions of the acid loop and the phosphate binding P-loop ((H/V)CX_5_R(S/T)) are highlighted in purple. Of the non-archaeal PTPs shown here, SHP-1 is an important anticancer drug target,^96^ and PTP1B and YopH are two of the most studied PTPs to date.^41, 91^ ShufPTP shows 90% sequence identity to TgPTP, 85% to TkPTP, 39% to SsoPTP, 25% to SHP-1, 30% to PTP1B and 31% to YopH, respectively.

Unsurprisingly, ShufPTP shows highest sequence similarity to the PTPs from *Thermococci*, the currently uncharacterized TgPTP, and TkPTP^56^ (90% and 85% sequence identity, respectively, see **Figures 3** and **S1,** with E-values of 8e-101 and 2e-91, sequence alignment was performed using the Standard Protein BLAST (BLASTp) webserver^92, 93^). This high sequence similarity is to be expected given both PTPs were used for sequence shuffling. As shown in **Figure S1**, not only is there high sequence similarity between ShufPTP and its progenitors, but also between the progenitors themselves: ShufPTP is separated by its closest progenitor, TgPTP by 15 amino acid substitutions, and the two most different progenitor proteins from each other, namely the PTPs from *T. celericrescens* and *T. gorganarius*, still share 75% sequence identity with each other. Given this, we consider ShufPTP to be a product of all five selected sequences.

Notably, as shown in **Figure 3**, the P-loops ((H/V)CX_5_R(S/T)) of the PTPs used in the sequence shuffling are identical, the acid loops (IPD-loop) of TgPTP and ShufPTP are identical, and the acid loops of TkPTP and ShufPTP differ only in one position (where a serine in ShufPTP presents as a glycine in TkPTP, likely increasing the flexibility of the TkPTP IPD-loop). While many PTPs have a WPD sequence in their acid loop, the archaeal PTPs shown in **Figure 3** carry instead an IPD sequence, which likely contributes to rigidifying the IPD-loop compared to the acid loops of PTPs such as PTP1B and YopH (available crystal structures of SsoPTP and TkPTP, PDB IDs: 7MPD,^55^ 2I6I,^94^ 2I6P^94^ and 5Z5A,^56^ 5Z59,^56^ all show the IPD-loop in its catalytically closed conformation). The P-loop GG motif shared among Archaea PTPs is thought to contribute to the unusual P-loop conformational flexibility that has been observed in TkPTP and SsoPTP.^55, 56^ However, mutational experiments on TkPTP have suggested that while the GG motif is important (but not sufficient) for P-loop flexibility, the GG-motif does not affect turnover.^56^

### Structural Characterization and Cysteine Oxidation

#### Structural Characterization

Four crystal structures of ShufPTP were determined ranging from 1.5 to 2.1 Å resolution (**Table S1**). In each case, ShufPTP adopts a canonical architecture characteristic of the PTP family, with particularly close structural similarity to its archaeal relatives TkPTP (RMSD 1.1 Å) and SsoPTP (RMSD 2.4 Å). The enzyme contains all four signature catalytic motifs that define modern PTPs: the phosphate-binding P-loop with conserved (H/V)CX_5_R(S/T) motif, the Q-loop that carries the water-directing glutamine, the IPD-loop that carries the conserved general acid (an aspartic acid), and the E-loop that carries a possible alternate general acid candidate (a glutamic acid).^97, 98^

In each of the ShufPTP structures, the Q-loop, IPD-loop, and E-loop conformations are largely superimposable, except for sidechain rotamer differences in E132 in the Q-loop and E38 and E39 of the E-loop (**Figure 4**). In each structure, the conventional general acid D63 on the IPD-loop adopts what appears to be a catalytically unproductive conformation, with its side chain orientated away from the P-loop, directed towards E41 on the E-loop. In structures in which the transition state analog vanadate was bound, it is coordinated by E132 and Q136 on the Q-loop. The interaction with Q136 is not typically observed in other vanadate-bound PTP structures and appears to be facilitated by an upward shift in the vanadate position relative to other PTP structures.

**Figure 4.**
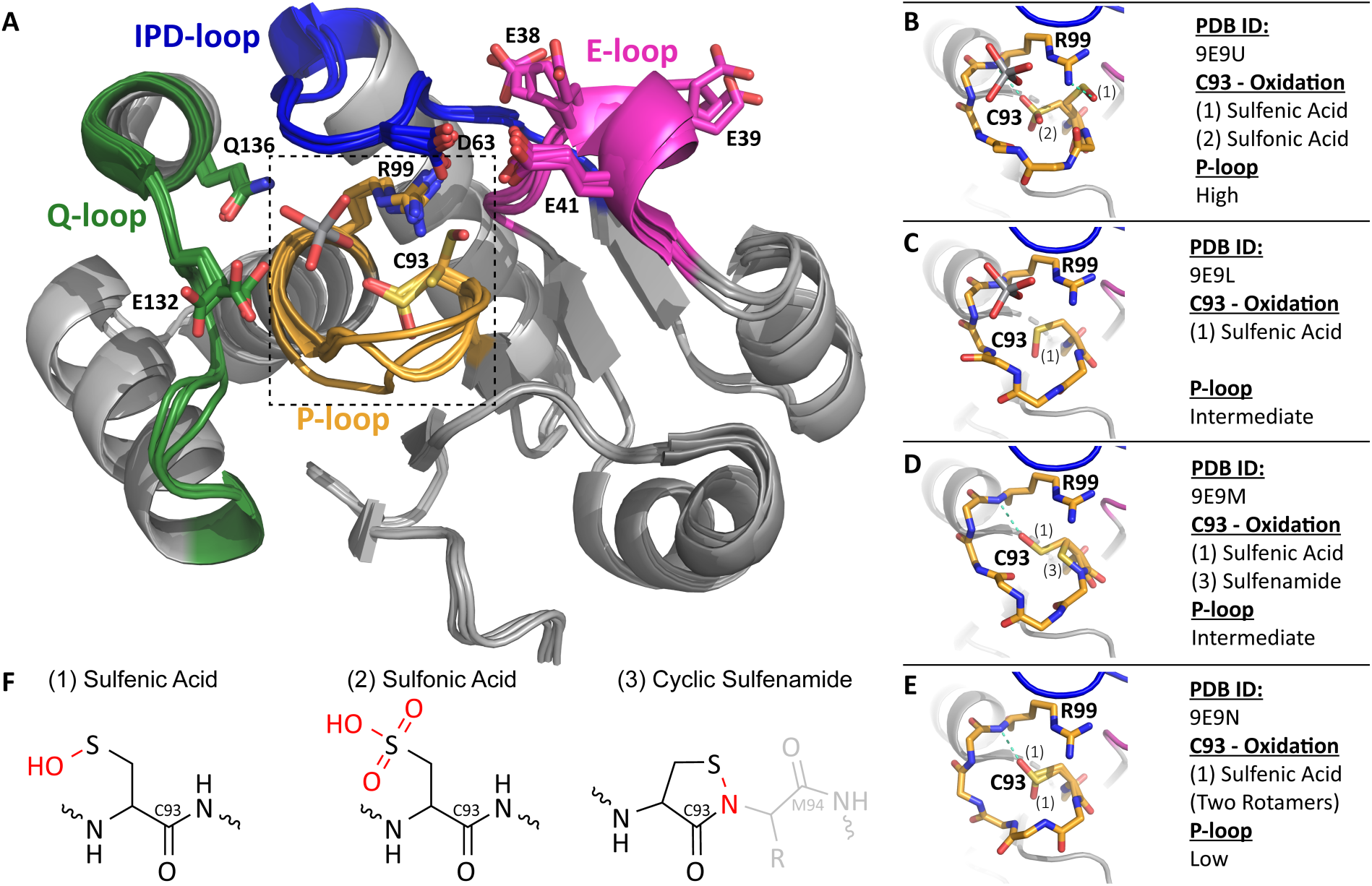
ShufPTP adopts a canonical PTP structure. (**A**) The active site of ShufPTP consists of four loop motifs, P-loop (Yellow), IPD-loop (blue), Q-loop (green), and E-loop (pink). All solved structures are largely superimposable with key differences seen in the P-loop motif. The vanadate ligand from PDB ID: 9E9U (this work) and key residues are shown in sticks. The dashed line refers to the P-loop region depicted in higher resolution in panels B-E. (**B**-**E**) Stick view of the different P-loop conformations and oxidation states. H-bonds are shown in teal, with differences in observed oxidation state, ligand, and conformation are listed to the right. (**F**) Chemical structure of the three observed oxidation states of C93. Atoms and bonds shown in red highlight the newly formed bonds caused by oxidation of the thiol.

The most striking difference between the ShufPTP structures is in the P-loop. We observe three distinct P-loop backbone conformations, which we designate “low,” “intermediate,” or “high,” referring to the relative position of the loop to the substrate binding site within the active site (**Figure 4**). The low conformation is identical to the catalytically competent form previously observed in TkPTP (PDB ID: 5Z5A).^56^

Remarkably, the oxidation states of the catalytic nucleophile, C93 differ depending on the structure, often with multiple oxidation states observed in a single structure (**Figures 4** and **S2**)). Specifically, we observe the cysteine as: (1) a sulfenic acid, (2) a sulfonic acid, and (3) a rare cyclic five-membered sulfenamide ring (**Figures 4B** through **F**). This latter species arises from an intra-molecular dehydration reaction between the sulfenic acid and an adjacent backbone amide from M94. In structures containing the sulfenic acid form, C93 does not form the canonical hydrogen bond to the central vanadium atom of the ligand, but instead is positioned away from the active site and in one case, forms a hydrogen bond with the conserved arginine R99 on the opposite side of the active site. We note that oxidation was observed in structures collected both on a home source and using synchrotron radiation. Thus, the observed oxidation appears to be a function of the susceptibility of ShufPTP to oxidation rather than the result of excessive X-ray radiation.

The relationship between the P-loop conformation and the oxidation state of C93 is unclear, in part because multiple oxidation states are often observed in a single structure (**Figure 4**). However, a general observation is that the oxidized C93 interacts with R99 in all conformational states except for the intermediate, vanadate bound conformation. C93 interacts with the sidechain of R99 only in the high conformation. In all cases, the R99 conformation is the same, regardless of P-loop conformation, ligand, or C93 interaction, suggesting that R99 could interact with vanadate in all P-loop conformations.

#### Enzyme Inactivation by Hydrogen Peroxide

Reversible oxidation of cysteine residues plays an essential regulatory role in enzyme catalysis and gene regulation,^99-103^ and specifically, oxidation regulates tyrosine phosphorylation.^104-107^ Several *in vitro* studies have identified hydrogen peroxide as a specific inactivator for PTPs.^70, 71, 98, 108-114^ The presence of the cysteine sulfenic acid and sulfonamide intermediate forms support the theory of reversible cellular redox-regulated PTP activities. In order to rationalize the observed oxidation states of C93 in the X-ray crystal structures (**Figure 4**), we measured the second order rate constant for inactivation of ShufPTP by hydrogen peroxide and compared it to reported values for extant PTPs (**Table 1**). ShufPTP exhibits a higher oxidation rate than any of the previously reported values for PTPs, with a *k*_inact_ of 65 ± 5 M^-1^ s^-1^.

**Table 1.** ShufPTP exhibits a higher rate of inactivation by hydrogen peroxide than other PTPs.

| Enzyme | $k_{\text{inact}}, \text{M}^{-1}\text{s}^{-1}$ |
| --- | --- |
| PTP1<sup>98</sup> | $9.1 \pm 0.1$ |
| VHR<sup>98</sup> | $17.9 \pm 1.3$ |
| LAR<sup>98</sup> | $14.0 \pm 3.1$ |
| YopH<sup>a</sup> | $33.0 \pm 4.1$ |
| PTP1B<sup>a</sup> | $18.5 \pm 1.6$ |
| ShufPTP<sup>a</sup> | $64.8 \pm 5.0$ |

<sup>a</sup> Measurements performed in this work.

#### Exploring Cysteine Oxidation in PTPs

Cysteine oxidation in proteins occurs as the result of an interplay between the protein electrostatic environment, active site solvation, redox potential, and the p*K*_a_ of the cysteine.^115^ Unfortunately, and particularly considering their high computational cost, microscopic approaches for computational prediction of cysteine p*K*_a_s, show poor performance in benchmarks, with p*K*_a_ root mean square error (RMSE) values > 3.5.^116^ Alternatively, the Gaussian-based method DelPhiPKa^117^ shows an improved RMSE of 1.7 in benchmarks, but is typically performed on a single structure at a time.^115^ This approach uses a smooth dielectric function to describe the dielectric properties of the system and mimic plausible conformational changes associated with changes in ionization states.

Despite the large errors in quantitative accuracy, DelPhiPKa can, however, predict cysteine p*K*_a_s with qualitative accuracy,^118^ making a relative comparison of the p*K*_a_ of the nucleophilic cysteine between different PTPs feasible. We performed qualitative comparison of the relative predicted p*K*_a_s of the nucleophilic cysteine based on extracting an ensemble of 50 evenly spaced snapshots from 5 x 500 ns molecular dynamics simulations of YopH (PDB ID: 2I42^43^), PTP1B (PDB ID: 6B90^119^), ShufPTP (PDB IDs: 9E9N and 9E9U, this work), and TkPTP (PDB ID: 5Z5A^56^ and 5Z59^56^). Given the conformational plasticity of the active site P-loop with cysteine oxidation state (**Figure 4**), we also included snapshots from simulations of ShufPTP and Tk-PTP initiated from X-ray crystal structures with their P-loops in their low/active and high/inactive conformations, to assess local dependence of the cysteine p*K*_a_ on the loop conformation. The resulting data, as well as corresponding experimental data for comparison, is shown in **Table S2**. As can be seen from this data, and from **Table 1**, the differences in *k*_inact_ by hydrogen peroxide for ShufPTP, PTP1B and YopH are small from a thermodynamic perspective, with only a 3.5-fold difference between the highest and lowest values. Therefore, unsurprisingly, our p*K*_a_ calculations are unable to capture this trend given the broader challenges with calculating cysteine p*K*_a_s discussed above. Nevertheless, our calculated p*K*_a_ values and thiolate fractions for the nucleophilic cysteine do qualitatively track with the measured *k*_cat_ values, where the thermodynamic differences are larger (**Tables 1** and **2**). Further, in the case of TkPTP and ShufPTP where crystal structures available for both the active/low and inactive/high conformations of the P-loop, we observe a slightly higher tendency for thiolate formation in the active/low P-loop states than in the inactive/high states.

**Table 2.** ShufPTP exhibits comparable turnover numbers to modern PTPs.^a^ ^a^ Data in Table 1 are *k*_cat_ values obtained at the respective pH optimum for each enzyme, at 25 °C except for YopH (22 °C) and TkPTP (20 °C). Data were obtained using the substrate *p*NPP except for TkPTP, where 6,8-difluoro-4-methylumbelliferyl phosphate (DiFMUP) was used. Note that *k*_cat_ reflects the second step of phosphoenzyme hydrolysis, which is substrate independent.

| Enzyme | $k_{\text{cat}}$, s<sup>-1</sup> | Reference |
| --- | --- | --- |
| YopH | 720 | 128 |
| PTP1B | 52 | 131 |
| VHZ | 3.9 | 82 |
| VHR | 3.1 | 82 |
| SsoPTP | 3.2 | 55 |
| TkPTP | 4.7 | 56 |
| TkPTP D63N | 0.01 | 56 |
| ShufPTP | 8.6 | This work |
| ShufPTP D63N | 0.12 | This work |

<sup>a</sup> Data in **Table 1** are $k_{\text{cat}}$ values obtained at the respective pH optimum for each enzyme, at 25 °C except for YopH (22 °C) and TkPTP (20 °C). Data were obtained using the substrate *p*NPP except for TkPTP, where 6,8-difluoro-4-methylumbelliferyl phosphate (DiFMUP) was used. Note that $k_{\text{cat}}$ reflects the second step of phosphoenzyme hydrolysis, which is substrate independent.

We further analyzed solvent accessibility across 5 x 500 ns MD simulation of different PTPs and the P-loop conformations, with the catalytic cysteine in each PTP in its deprotonated form. Histogram and solvent density analyses of these simulations (**Figure 5**) showed low solvent stabilization of the nucleophilic cysteine in PTP1B, slightly greater stabilization in YopH, and a clearly increased solvent density in Tk-PTP and ShufPTP, which mirrors the trend in *k*_inact_ by hydrogen peroxide across these systems (**Table 1**). Of the systems studied, only ShufPTP and Tk-PTP were able to simultaneously stabilize 3 water molecules within a 5Å sphere centered on the S_γ_-atom of the catalytic cysteine, creating greater possibility for cysteine oxidation to occur than in PTP1B and YopH.

**Figure 5.**
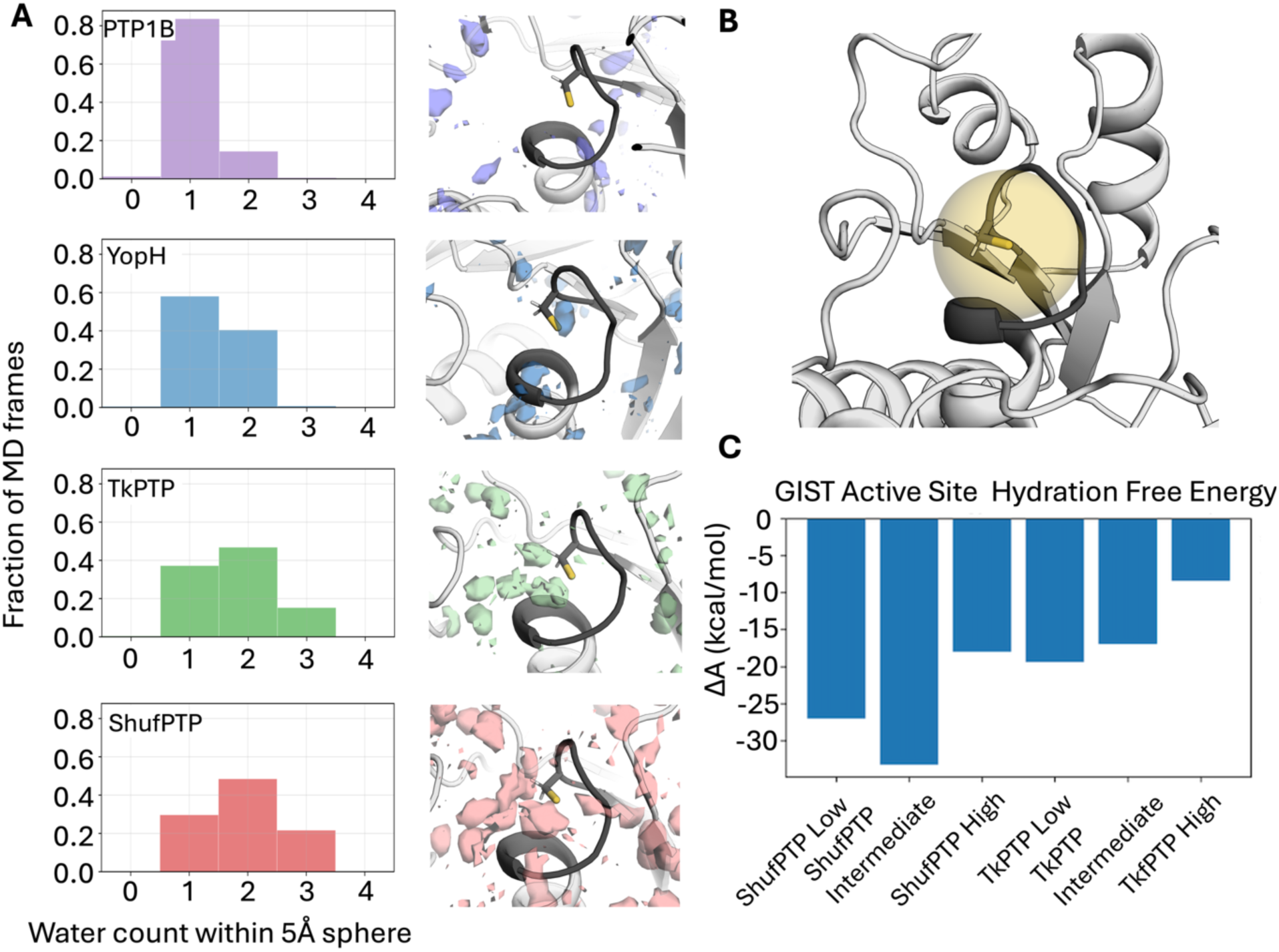
Comparison of active site solvent penetration across human and archaeal PTPs considered in this work. **(A)** Water density and a histogram analysis of water molecules present within a 5Å sphere centered on the S_γ_-atoms of the nucleophilic cysteine sidechains of PTP1B (PDB ID: 6B90^119^), YopH (PDB ID: 2I42^43^), TkPTP (PDB ID: 5Z5A^81^), and ShufPTP (PDB ID: 9E9N, this work), in the low/active P-loop and closed acid loop states (in the case of ShufPTP). Analysis was based on 5 x 500ns MD simulations per system. **(B)** Visualization of the 5Å sphere centered around the nucleophilic cysteine S_γ_-atom used for evaluation of active site solvation**. (C)** Free energy of solvation computed according to Grid Inhomogeneous Solvation Theory,^120^ and integrated within the 5Å analysis sphere shown in panel **B**. Evaluation was performed of solvent penetration in the low/active, intermediate and high/inactive states of ShufPTP and TkPTP. Standard deviations across three replicas are less than 0.3 kcal/mol and were thus not included for clarity. The raw data for this figure is provided in the associated Zenodo data submission, DOI: 10.5281/zenodo.15074903.

Furthermore, the role of P-loop dynamics in solvent stabilization of the active site was assessed *via* Grid Inhomogeneous Solvation Theory (GIST)^120^ analysis on restricted simulations initialized from structures that capture high/inactive, intermediate, and low/active states of the P-loops in ShufPTP and TkPTP. The active site hydration free energy was computed using the standard GIST protocol, and integrated within the 5Å region surrounding the S_γ_-atom of C93.^121^ The high/inactive P-loop states of both enzymes was characterized by the lowest free energy of solvation, which supports the p*K*_a_ calculations shown in **Table S2**. Interestingly, the P-loop active (low) and intermediate states of ShufPTP showed free energies of solvation 7.6 and 16.3 kcal/mol lower than observed in TkPTP. The increased solvation of the ShufPTP active site compared to counterparts can account for the correspondingly more facile oxidation of the active site cysteine nucleophile.

### ShufPTP Substrate Preference, Mechanism and Catalytic Backups

#### Substrate Preference by Naphthyl Phosphate Isomers

The β/α-naphthyl phosphate hydrolysis ratio serves as a reporter for the active site geometry and substrate preferences for PTPs.^97, 122-126^ The PTP superfamily includes enzymes specific for phosphorylated tyrosine residues, and the dual-specificity phosphatases (DUSPs) that hydrolyze phosphorylated serine and threonine residues as well as tyrosine.^91, 127^ The substrate specificity of tyrosine-specific PTPs arises from a deeper and narrower catalytic site pocket compared to those of DUSPs. Although the β/α-naphthyl phosphate hydrolysis ratio has not been measured for the five progenitor PTPs used for the sequence shuffling (**Figure S1**), it has been measured for six other PTPs, and shows clear differentiation between DUSPs and classical PTPs.^115^ Specifically, due to the steric and structural resemblance of α-naphthyl phosphate to phosphoserine and phosphothreonine, and β-naphthyl phosphate to phosphotyrosine, extant DUSPs exhibit a β-NP/α-NP (*V/K* ratio) in the range of 1 – 2, compared to a ratio of 7 or higher for classical tyrosine specific PTPs.^115^ Furthermore, given the P-loop conformational changes of the similar TkPTP due to heat treatment,^56^ it was suspected that ShufPTP might exhibit similar temperature-dependent conformational changes, and thus, a change of active site geometry and substrate preference. The ratio of (*V/K*) for β- to α-naphthyl phosphate substrate consumption catalyzed by ShufPTP is 1.7 at 22 °C and 1.8 at 60 °C, indicating that ShufPTP does not exhibit temperature-dependent changes in active site geometry, which is more analogous to extant members of the dual-specificity phosphatases (DUSP) subfamily of PTPs, than tyrosine-specific classical PTPs.

#### Phosphatase Activities and Dual General Acid Catalysis

ShufPTP exhibits a bell-shaped pH-rate profile like extant PTPs (**Figure 6)**, indicating a conserved catalytic mechanism involving two essential catalytic residues (**Figure 1**). The turnover number for ShufPTP is 8.6 ± 0.1 s^-1^ at its pH optimum 4.75 at 22 °C, approximately two-fold faster than its closest extant relative, TkPTP.^56^ **Table 2** compares *k_cat_* values of ShufPTP to extant PTPs. The *K_M_* for the substrate *p*NPP at ShufPTP’s pH optimum is 2.3 mM, within the range of other PTPs at their respective pH optima (YopH: 2.55 mM;^98^ VHR: 1.59 mM;^128^ Sso: 3.4 mM;^55^ VHZ, 8.3 mM).^97^

**Figure 6.**
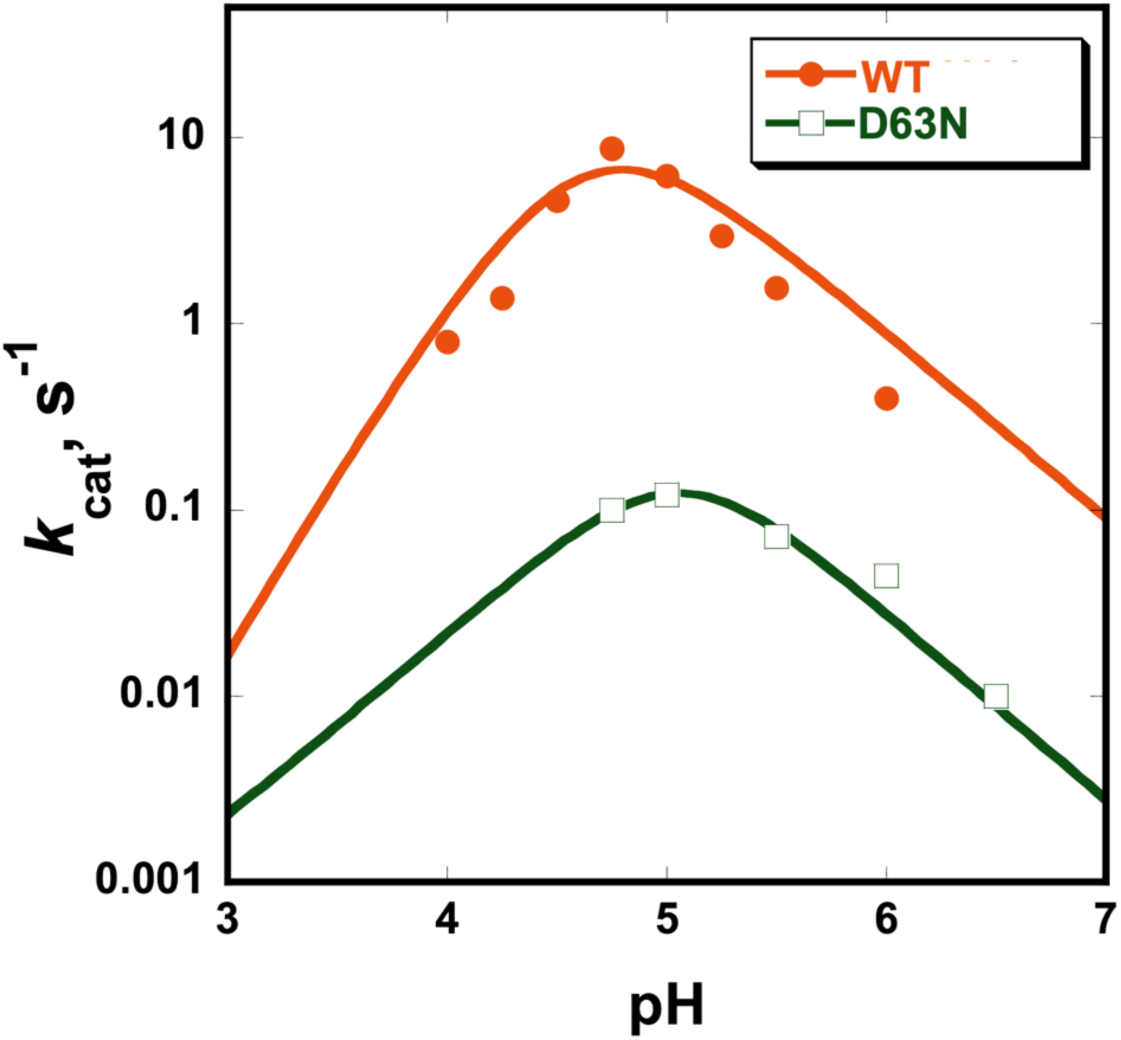
ShufPTP exhibits a bell-shaped pH-rate profile like extant PTPs. The retention of the basic limb in the profile for the general acid variant D63N indicates the presence of an alternate general acid. Precipitation of the variant at pH 4.5 or below precluded acquisition of data at lower pH. The plot was thus fit speculatively to a Bell equation, as the Cysteine nucleophile must still be present and active, otherwise there would be no activity. The presence of the basic limb, however, clearly shows the presence of general-base catalysis also in the D63N variant.

By analogy with the structures of extant PTPs, residue Asp63 in ShufPTP occupies the position of the conventional general acid. The ShufPTP variant D63N has a turnover number of 0.1 s^-1^ at pH 5.0, approximately 10-fold faster than the corresponding variant of TkPTP, which is also D63N^56^ (**Table 2**). Below pH 4.5, the variant exhibited precipitation, attributed to denaturation, limiting the collection of kinetic data on the acidic limb of the pH-rate profile. Thus, the fit of the curve to a Bell-equation on the acid limb is speculative. However, it is worth noting that although the decrease between pH 5.0 and 4. seems visually small, this plot is on a log scale and thus the changes are much larger than they visually appear. Further, the presence of a clear basic limb shows that also the D63N variant uses general acid/base catalysis. Notably, this general acid variant of ShufPTP exhibits an approximately 70-fold rate reduction compared to the native enzyme, and the basic limb of the pH-rate profile is retained. This contrasts with the typical kinetic characteristics of the general acid D to N variants of PTPs, where a 2 to 3-order of magnitude lower activity and loss of the basic limb of their pH-rate profiles are observed.^129, 130^ A small number of characterized PTPs exhibit the phenomenon of a smaller than expected rate reduction and the retention of a bell-shaped pH profile in their general acid variants, a combination shown to result from the presence of a second acidic residue in or near the active site that than can function as an alternate general acid, as documented in VHZ^82^ and SsoPTP.^55^

#### Identifying Prospective Catalytic Backup Residues

In order to identify the prospective backup acid, we created D63N/E41Q and D63N/E132Q variants of ShufPTP, thus knocking out both prospective backup acids. Neither variant showed activity, however, and both variants expressed poorly and had different chromatographic behavior compared to the native enzyme and the D63N variant (which behaved similarly to one another). We presume that this is due to conformational changes induced by the double mutations; however, this precludes being able to identify the backup residue through mutation. Thus, we have thus instead performed a detailed computational analysis, as outlined below.

Visual examination of all available crystal structures of ShufPTP led to the identification of several carboxylate side chains located near the active site (**Figure S3**). While some of these residues point away from the active site, they are all located on mobile loops (D63 on the IPD-loop, E38, E39 and E41 on the E-loop, and E132 on the Q-loop), making it plausible that loop fluctuations could render these side chains transiently conformationally accessible for catalysis, allowing them to act as backups.

To validate this hypothesis, we tracked the distances between the side chains of E38, E39, D63, E41 and E132 during MD simulations of the phosphoenzyme intermediate states of low P-loop state wild-type and D63N ShufPTP (PDB:9E9N, this work), at both 300 and 360K (**Figures 7**, and **S4**). For this analysis, we focused our simulations on the phosphoenzyme intermediate state of ShufPTP, as we expect hydrolysis to be rate-limiting, based on burst kinetics of PTP1B and YopH.^45, 132^ The low structure of the P-loop was chosen for this analysis, due to its structural similarity to the catalytically active state of TkPTP.^56^ Additionally, when simulated in the absence of the phosphate, this starting structure was the only one out of four ShufPTP crystal structures that demonstrated positional stability of the side chain R99, which is a key conserved residue important for phosphate binding in the active site^133^ (**Figure S5**).

**Figure 7.**
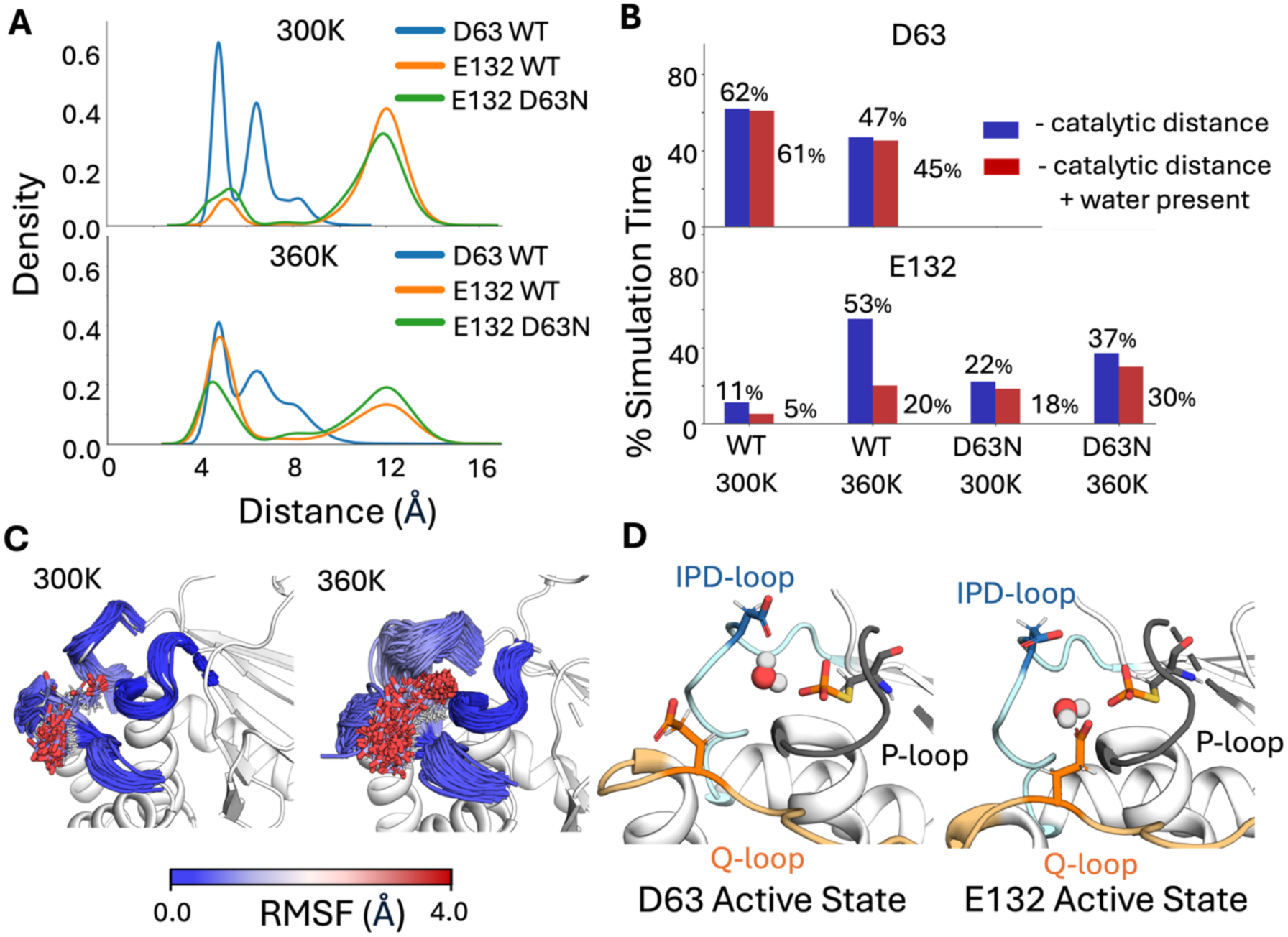
Characterizing prospective catalytic backups in ShufPTP. Potential backups identified by tracking the distances between the phosphate group at the phosphoenzyme intermediate and prospective catalytic carboxylate side chains, extracted from our MD simulations of wild-type and D63N ShufPTP. (**A**) Kernal density estimates (KDE) of the distances between the phosphate group and prospective catalytic residues, defined as the distance between the P atom of the phosphate group and the closest oxygen atom of the carboxylate side chain of each key residue. (**B**) The % simulation time the side chains of each of the three most likely candidates (D63, E41 and E132) spent within 6Å of the phosphate group phosphorus atom (shown in blue). Further, to be catalytically viable, it is necessary for a water molecule to bridge the respective side chain and the phosphate group, in order to act as a nucleophile (Figure 1). For each prospective catalytic side chain, we calculated the % of simulation time a water molecule is positioned within both 3.5Å of the carboxylic acid of the respective side chain, and of the phosphorus atom of the phosphate group (red, measured based on distances to the nucleophilic oxygen atom). (**C**) The dramatic increase in proximity of E132 to the C93 side chain with increased temperature can be attributed to the increased flexibility of the Q-loop. (**D**) Illustration of the catalytically active conformations of D63 (IPD-loop) and E132 (Q-loop), with a catalytic water molecule bridging the side chain and the phosphate group.

Based on the analysis of phosphorylated wild-type and D63N simulations, only two residues sample conformations within a plausible reactive distance of the phosphoenzyme intermediate in the wild-type enzyme: the native acid, D63, and E132, which is located on the Q-loop. The distance distribution of E132 shows two peaks: a dominant non-reactive peak, and a second, smaller peak at ∼4.1 Å, which plausibly leads to a reactive conformation. We repeated this analysis at 300K in the D63N variant, and at 360K in both variants (**Figures 7** and **S4**), demonstrating that both abolishing the native catalytic residue and increasing temperature increases sampling of prospectively catalytic conformations of E132, during the simulations of both ShufPTP and of the D63N variant.

We note that for the hydrolysis reaction to occur, it is essential to have a water molecule correctly positioned for both activation by the respective glutamic acid side chain, and for nucleophilic attack on the phosphorus group of the intermediate. To account for this, we also tracked water molecules that bridge the D63, E132, and E41 side chains and the phosphate group, and are correctly positioned for reaction (*i.e.* within 3.5Å of both the carboxylate and phosphate groups) during our simulations (**Figures 7B and S4B**), indicating that E132 is at least transiently aligned for catalysis during our simulations, in particular at 360K (where we see a drop in catalytic conformations of D63). Finally, as shown in **Figure 7C** and **D**, the increase in catalytic conformations of the E132 side chain at higher temperature are facilitated by increased flexibility of the Q-loop, allowing for a coordinated rearrangement of the IPD- and Q-loops. This rearrangement could facilitate the role of E132 as a backup.

Finally, E132 replaces one of two typically invariant glutamine residues on the Q-loop, which are essential for nucleophile positioning in the second hydrolysis step of the PTP-catalyzed reaction (**Figure 1**).^134^ In doing so, ShufPTP introduces a residue potentially capable of acid-base catalysis into this loop. Importantly, this Q-loop substitution is not unique to ShufPTP, and is also observed in Tk-PTP, and VHZ, both of which have been suggested to use back-up catalytic mechanisms ^56, 82^ (**Figure S6**). This substitution is not observed in the archaeal PTP SsoPTP, which instead achieves a catalytic backup mechanism through using a glutamic acid (E40) in an analogous position to E41 in ShufPTP.^55^ This significantly increases the likelihood that this single amino acid substitution is sufficient to also impact backup catalytic activity to ShufPTP, as explored by empirical valence bond simulations in the subsequent section.

#### Empirical Valence Bond Simulations of Phosphoenzyme-Intermediate Hydrolysis

To verify whether our hypothesis with regards to the role of E132 as a catalytic backup is correct, we performed empirical valence bond (EVB) simulations of phosphoenzyme-intermediate hydrolysis in wild-type and D63N ShufPTP, facilitated by either D63 or E132 in each enzyme. Based on prior work on other enzymes with catalytic backups, one would expect that any backup mechanism observed in the D63N variant would also exist in the wild-type enzyme, albeit less effectively than the native mechanism.^135-137^ Because the crystal structure of ShufPTP demonstrates only catalytically unproductive conformation of the D63 side chain (with D63 pointing outside the active site and forming what appears to be a low-barrier hydrogen bond with the side chain of E41 on the E-loop), EVB simulations were initialized from snapshots taken from our MD simulations with catalytically active Asp/Glu conformations, as described in the **Materials and Methods**.

We note that for our EVB simulations, as in our MD simulations, we focus on modeling the hydrolysis of the phosphoenzyme intermediate rather than nucleophilic attack on the Michaelis complex (**Figure 1**), as hydrolysis of this intermediate is expected to be the rate limiting step of catalysis.^45, 132^ Beyond investigating the impact of the choice of the residue that facilitates the reaction, we assessed whether the complementary proximity of D63 and E132 to the active site has a noticeable effect on the activation barrier of the reaction (*i.e.* whether both or only one of these side chains points into the active site simultaneously). Therefore, to represent a complete set of plausible electronic environments, we considered the five following catalytic scenarios: (1, 2) wild-type ShufPTP system with D63 as the catalytic residue and E132 pointing in or out of the active site, (3, 4) wild-type ShufPTP with E132 as the catalytic residue and D63 pointing in or out of the active site, and (5) the D63N ShufPTP mutant, in which the reaction is facilitated solely by E132 (**Figures 8** and **S7,** and **Table S3)**.

**Figure 8.**
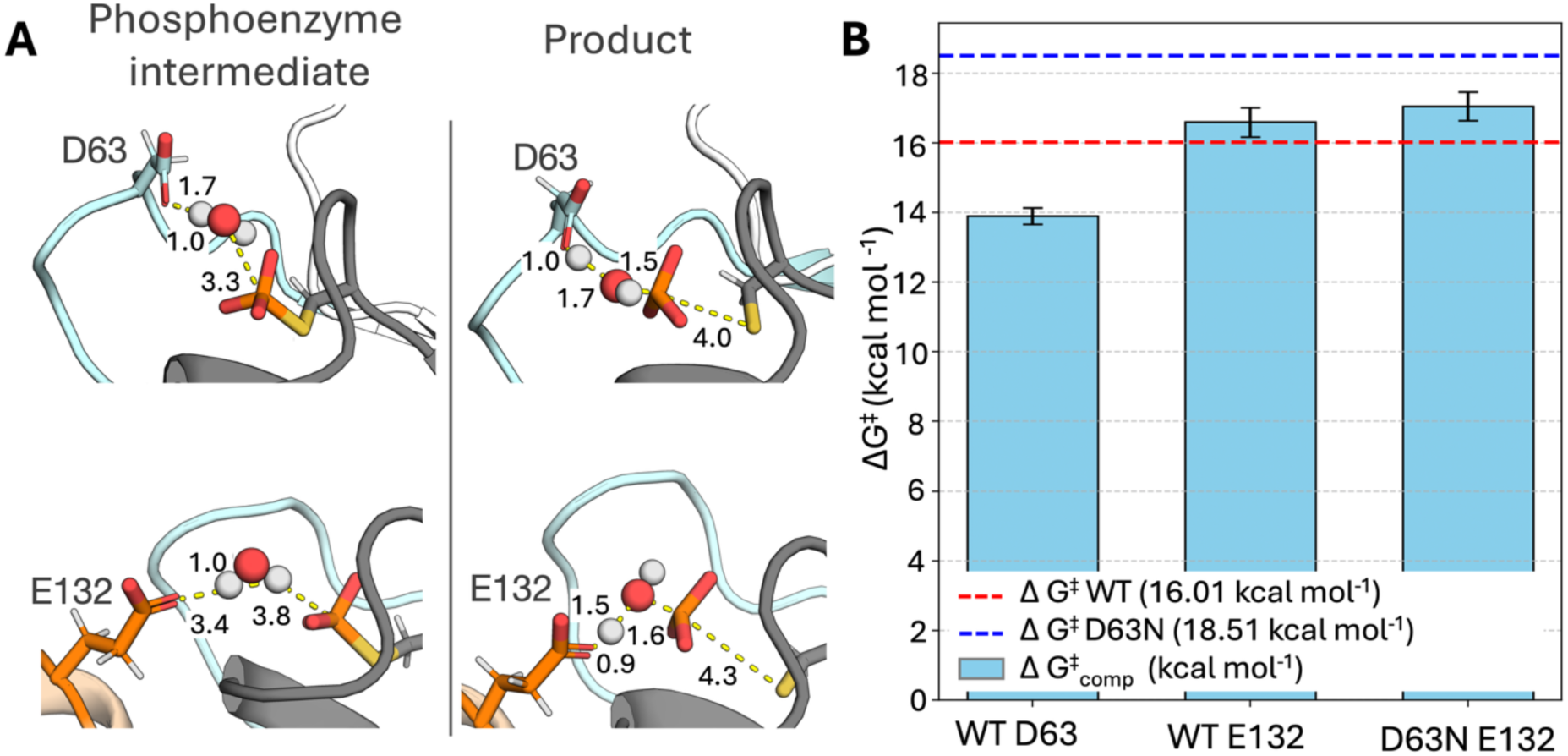
Empirical Valence Bond evaluation of reaction barrier for the dominant and backup mechanism of ShufPTP. **(A)** Representative snapshots of phosphoenzyme intermediate and product state structures extracted from empirical valence bond (EVB) simulations^138^ of the hydrolysis reaction catalyzed by wild-type ShufPTP. Shown here are stationary points for the D63-as-base and E132-as-base mechanisms. (**B**) Calculated ΔG^‡^ values (kcal mol^-1^) for the D63-as-base and E132-as-base mechanisms in wild-type (WT) and D63N ShufPTP were compared to the experimental values obtained from the kinetic data (*k*_cat_, **Table 2**) for both variants. The red dashed line indicates the experimental activation free energy for the reaction catalyzed by wild-type (WT) ShufPTP, and the blue dashed line indicates the experimental activation free energy for the reaction catalyzed by the D63N ShufPTP mutant. The error bars represent the standard error of the mean on the calculated activation free energies over 20 individual EVB trajectories for each system. The raw data for this figure are shown in **Table S3**.

Our EVB simulations (performed at 300K) indicate that the energetically preferred active site conformation for ShufPTP is one where both the D63 and E132 side chains point into the active site, as the electrostatic repulsion between these two side chains facilitates easier proton abstraction from the nucleophilic water molecule (**Figure 1**). In wild-type ShufPTP, in all scenarios, proton abstraction in a D63-as-base mechanism has a lower activation free energy than in an E132-as-base mechanism. However, the difference in energy between the two mechanisms in the preferred conformation (both side chains pointing into the active site) is only 2.7 kcal mol^-1^, with a calculated activation free energy of 13.9 kcal mol^-1^ for the D63-as-base mechanism, and a calculated activation free energy of 16.6 kcal mol^-1^ for the E132-as-base mechanism. For reference, the corresponding experimental value is 16.6 kcal mol^-1^ (**Table S3**), based on a *k*_cat_ of 8.6 s^-1^ (**Table 2**), calculated using transition state theory. Thus, both plausible mechanisms are energetically within range of the corresponding experimental activation free energy, with a preference for the native D63-as-base mechanism.

The calculated activation free energies for E132-as-base mechanism in wild-type ShufPTP, obtained from starting conformations in which D63 points out of the active site (17.4 kcal mol^-1^, **Table S3**), are very similar to the calculated activation free energy for the D63N mutant (17.1 kcal mol^-1^). This suggests that (1) both D63- and E132-as-base mechanisms are catalytically plausible in wild-type ShufPTP, with D63 dominating due to both a lower activation free energy and more frequent sampling of catalytic conformations, and (2) at least part of the loss of activity in the D63N mutant is not due to the loss of catalytically favorable ground state destabilization, but rather due to elimination of the charge on the D63 side chain from the active site, making the E132-as-base mechanism about 0.4 kcal mol^-1^ (from EVB calculations) energetically less favorable than in wild-type ShufPTP.

### ShufPTP Thermostability

#### Experimental Characterization of ShufPTP Thermostability

To assess thermal stability, catalysis by ShufPTP was measured from 22 – 90 °C. ShufPTP exhibits a turnover number of 26 ± 3 s^-1^ at 60 °C, which is comparable to TkPTP at the same temperature.^56^ The fastest activity for ShufPTP was observed at 90 °C, with a turnover number of 140 ± 20 s^-1^, approximately 16-fold faster than that at 22 °C. Enzyme activity determinations for ShufPTP within the 22-90 °C temperature range show an increase in the steady-state rate with temperature that conforms to the Arrhenius equation (**Figure S8**). Protein denaturation within the studied temperature range would have brought about a decrease in activity with concomitant deviation from the Arrhenius behavior, which is not observed in the experimental data. The *K*_M_ for the substrate pNPP is 2.3 ± 0.4 mM at 22 °C and 1.9 ± 0.6 mM at 90 °C, further supporting retention of structural integrity in this temperature range. We conclude, therefore, that the thermal denaturation of ShufPTP takes place at temperatures significantly above 90 °C. Verification of denaturation at temperatures above 90 °C is challenging, because the 100 °C boiling point of water which sets an upper limit to the temperature range that can be accessed by many experimental methodologies. This constraint is overcome when using differential scanning calorimetry (DSC) to study denaturation, because DSC experiments customarily involve the application of a moderate overpressure on the order of a couple of atmospheres. While the purpose of this procedure was originally to prevent bubble formation during the temperature scan, the most relevant consequence of the applied overpressure is to increase the boiling point of water and enable the temperature scans to terminate many degrees above 100 °C.

We performed DSC experiments with solutions of ShufPTP at pH 7, under an overpressure of about 2 atm. Several DSC experiments were performed terminating at different temperatures, including one terminating above 120 °C. Remarkably, none of these experiments (see **Figure 9**) revealed a calorimetric transition (*i.e.,* a heat capacity “peak”) that can be attributed to protein denaturation. It is important to note that the absence of a calorimetric transition in the scans with ShufPTP cannot be attributed to instrumental limitations or to a low denaturation heat capacity signal for this protein. This is because the area under a calorimetric transition is the denaturation enthalpy, which as a first approximation, scales with the protein molecular mass.^139^ For a protein of a molecular mass of 17 kDa, such as ShufPTP, a simple calculation based on the known structure-energetics correlations (Table 5 in ref. ^139^) leads to estimates of the denaturation enthalpy of about 191, 215, 239 and 262 kcal/mol at 90 °C, 100 °C, 110 °C and 120 °C, respectively. These high denaturation enthalpies suggest that if the protein denaturation took place within the 90-125 °C temperature range, it would have been easily detected in our DSC experiments. In order to illustrate this, we performed DSC experiments with thioredoxin under the same conditions (in particular, using the same protein concentration as in the ShufPTP experiments: 1mg/mL). Thioredoxin has a molecular mass of about 11 kDa and, therefore, its denaturation enthalpy values, as estimated from the structure-energetics correlation, are substantially lower than those predicted for ShufPTP. Yet, the denaturation transition is clearly seen at about 90 °C in DSC scans with thioredoxin solutions of concentration 1 mg/mL.

**Figure 9.**
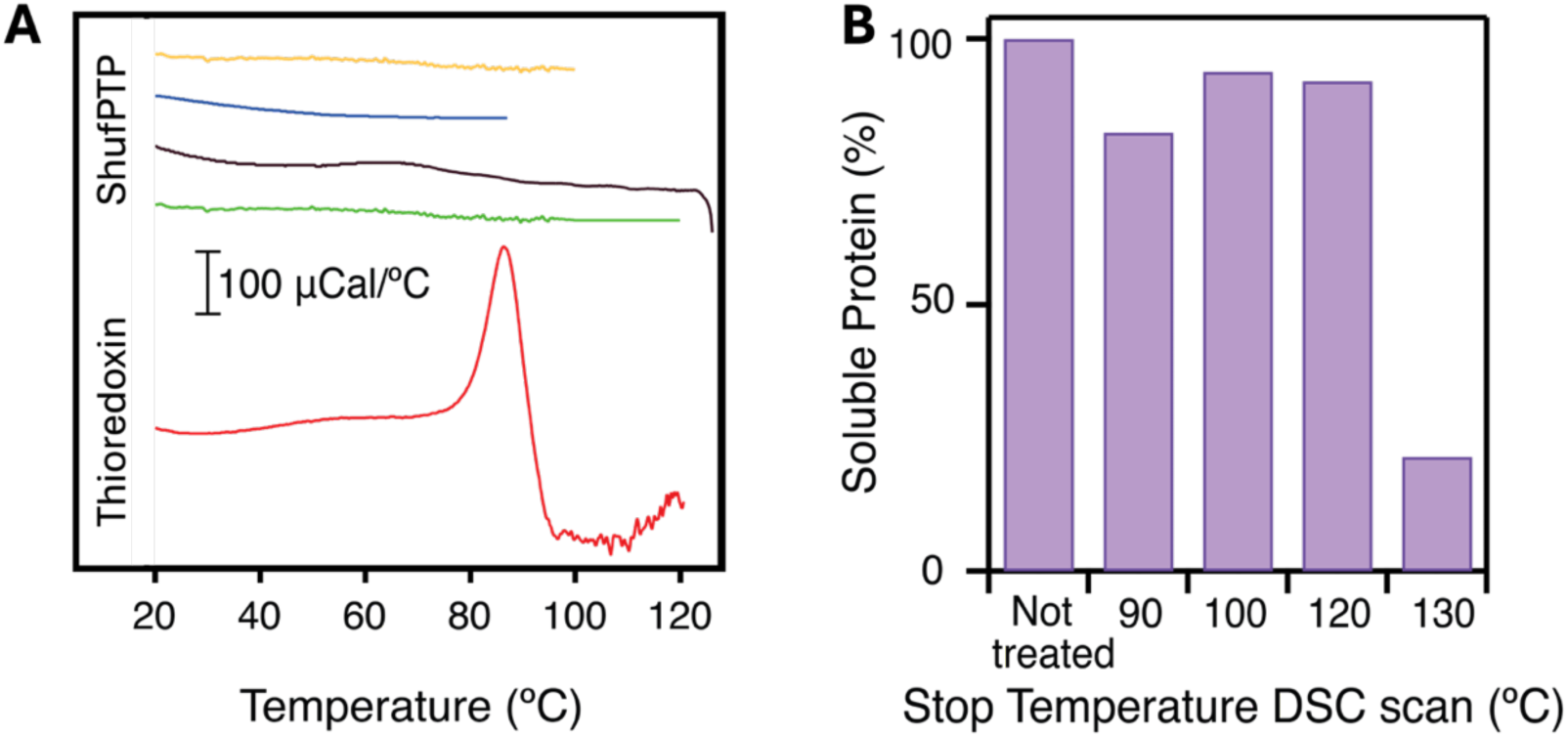
Differential scanning calorimetry (DSC) of ShufPTP. (**A**) Profiles of heat capacity versus temperature for solutions of ShufPTP and thioredoxin at 1 mg/mL and pH 7. The profiles have been shifted in the y-axis for the sake of clarity. The four uppermost profiles correspond to ShufPTP solutions and differ in the upper temperature of the DSC scan, as it is visually apparent. The lowermost profile corresponds to a thioredoxin solution and, unlike the profiles for ShufPTP, shows a prominent denaturation transition (heat capacity peak). (**B**) Amount of non-aggregated (soluble) protein in solutions of ShufPTP extracted from the calorimetric cell after cooling from a DSC scan. The amount of soluble protein is given as function of the highest temperature reached in the DSC scan.

Overall, the absence of a calorimetric transition in our DSC scans of ShufPTP solutions support a very high denaturation temperature for this protein. We found indirect evidence from the calorimetric experiments of the onset of denaturation at about 130°C. Thus, we detected substantial aggregation in the sample extracted from the calorimetric cell after cooling from a DSC scan in which a temperature of about 130 °C had been reached. On the other hand, aggregation was not observed in protein solutions extracted after cooling in experiments with a final temperature lower than 130 °C. For all these experiments, the protein solution extracted upon cooling was centrifuged to eliminate any aggregated protein, and the supernatant was subjected to gel electrophoresis. In all cases, a single band corresponding to the molecular weight of ShufPTP was observed, but its intensity was much smaller for the sample that had reached 130 °C in the calorimetric experiment, reflecting strong aggregation in this case (see lower panel in **Figure 9**).

These experiments can be rationalized in terms of the well-known Lumry-Eyring mechanism of irreversible protein denaturation,^140-142^ in which reversible protein unfolding is followed by an irreversible alteration of the unfolded protein to lead to an irreversible denatured state unable to fold back to the native protein upon cooling. Aggregation, as it is observed at about 130 °C for ShufPTP, is the most common process responsible for irreversibility in protein thermal denaturation. Therefore, the experiments reported in **Figure 9B** constitute in fact a reversibility test and the results obtained could in principle correspond to two different scenarios in the context of the two-stage Lumry-Eyring mechanism. Reversible unfolding could occur at temperatures substantially lower than the onset of irreversibility seen at about 130 °C. However, this scenario is immediately ruled out by the fact that we did not observe the unfolding transition in our calorimetric experiments. The second, and far more plausible, scenario is that unfolding and irreversible alterations occur in the same temperature range, implying that each unfolded molecule that is formed is immediately transformed in the irreversibly denatured (aggregated) state. It follows that the denaturation temperature higher than 130 °C predicted by our analyses would correspond to the global denaturation of the protein, although denaturation would correspond to a two-state irreversible denaturation, as is in fact the case for many proteins.^143^

#### Computational Characterization of ShufPTP Thermostability

Having established that ShufPTP has substantively higher thermostability than would be expected from any of its progenitor enzymes, the next question is *why* ShufPTP is so thermostable. While the high hydrophobic content of ShufPTP certainly plays a part^144-147^ (44.7% on the Kyte-Doolittle scale,^148^ compared to 41-44% for its progenitors and ∼35% for PTP1B and SHP-1), this is likely not the only contributing factor, as otherwise ShufPTP should behave similarly to TgPTP at 44% hydrophobic content. In fact, detailed analysis by Nussinov and colleagues^149^ shows several other factors as also playing important roles in facilitating protein thermostability, including the prevalence of increased salt bridges and side chain-side chain hydrogen bonds, increased frequency of Arg and Tyr side chains, reduced frequency of Cys and Ser side chains, and a larger fraction of residues in α-helical conformations, compared to their mesophilic counterparts. Further, statistical analysis has indicated that thermophilic globular proteins have a significantly higher aliphatic index (volume occupied by their aliphatic side chains) than mesophilic proteins.^150^

To explore the role of these features in facilitating the extreme thermostability of ShufPTP, we have analyzed various physical and chemical properties of a range of thermophilic archaeal PTPs, as well as mesophilic human and bacterial PTPs, using the ProtParam tool on the Expasy web-server.^151^ These include all five progenitor archaeal PTPs shown in **Figure S1**, additional archaeal PTPs TkPTP and SsoPTP, the human PTPs PTP1B, SHP-1 and TCPTP (T-cell Protein Tyrosine Phosphatase), the bacterial PTP YopH, and the atypical human dual-specificity phosphatase VHZ. We have specifically focused on comparing the following properties between enzymes: (1) the amino acid composition, (2) the instability index (providing a measure of the stability of a protein in a test tube, values <40 indicate stability and >40 stability ^152^), (3) the aliphatic index and (4) the GRAVY index (the Grand Average of Hydropathy, with more negative values being associated with greater hydrophilicity). Based on our analysis, we observe that, consistent with analysis by Nussinov and colleagues,^153^ we observe slightly more frequent occurrence of Arg and less frequent occurrence of Cys and Ser in the thermophilic archaea, as well as additionally slightly lower occurrence of Gln and higher occurrence of Glu and Leu side chains in the thermophiles compared to their mesophilic counterparts. We also see higher instability indexes and aliphatic scores in the thermophiles, and generally less negative GRAVY indexes. The exception to this is VHZ: similarly to the archaeal PTPs, TkPTP, SsoPTP and ShufPTP, VHZ has also been shown to function through a catalytic backup mechanism.^55, 56, 82^ Our sequence analysis shows that this atypical phosphatase has similar Cys, Gln and Glu composition and a similar instability index to its human and bacterial mesophilic counterparts, but a similar Arg and Leu composition and similar aliphatic and GRAVY indexes to the archaeal PTPs, placing it as a hybrid between the groups.

Importantly, while our sequence-based data show clear differences between thermophilic and mesophilic PTPs as a group, they do *not* allow us to differentiate enzymes within the groups, and thus provide insight into the thermostability of ShufPTP only in general terms. To further explore the thermostability of ShufPTP, we compared the prevalence of non-covalent interactions and the stability of helical features in 5 x 500 ns MD simulations of PTP1B, YopH, TkPTP and ShufPTP (**Figure 10**). The abundance of hydrophobic interactions, salt bridges and side chain hydrogen bonds was quantified in terms of the % residues engaged in the different interaction types, to account for variability in system size between PTPs.

**Figure 10.**
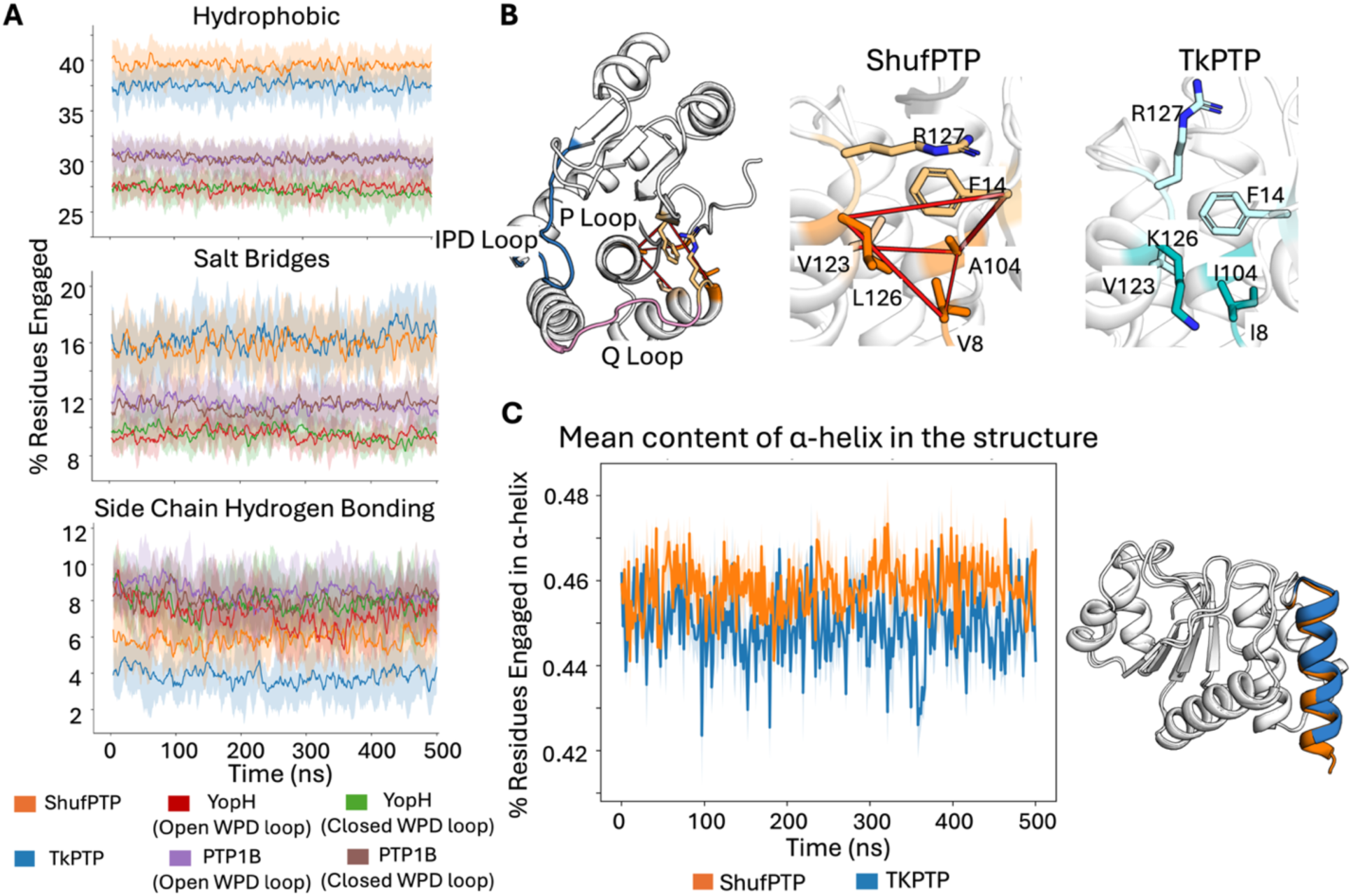
Computational analysis of structural properties potentially linked to ShufPTP thermal stability. Data shows analyses across 5 x 500 ns conventional MD simulations of each of TkPTP (PDB ID: 5Z5A^56^), ShufPTP (PDB ID: 9E9N, this work), PTP1B (PDB ID: 6B90^119^) and YopH (PDB IDs: 2I42^43^ and 1YPT^42^). Shown here are (**A)** the time evolution of non-covalent interactions across the trajectories for each system. Interactions have been quantified based on % residues engaged in the specific interaction type, to adjust for variations in protein size among the different systems. The mean value across all replicates for each system is highlighted in bold. **(B)** Representative examples of hydrophobic interactions uniquely present in ShufPTP and not in TkPTP, projected onto the structure of the respective PTP, to demonstrate the proximity of these interactions to the corresponding PTP active sites and to residues located on dynamic loops. **(C)** Time evolution of the mean content of residues engaged in an α-helix across replicates for each system.

As seen from these data, both thermostable ShufPTP and TkPTP demonstrate a higher prevalence of hydrophobic interactions and salt bridges during our simulations than PTP1B and YopH, which are more similar to each other. Additionally, ShufPTP shows a higher prevalence of side chain hydrogen bonding interactions than TkPTP (**Figure 10A**). While the differences between TkPTP and ShufPTP may seem visually negligible, the interactions that differ between the two PTPs often occur in strategic regions of the scaffold that engage typically dynamic protein regions. To illustrate this, **Figure 10B** shows an example of hydrophobic interactions that are uniquely present in ShufPTP and not in TkPTP. This set of residue substitutions engage a hydrophobic network located in the proximity of the active site, and allow for a cation-π interaction to be formed between the side chains of R127 of the Q-loop and F14 positioned on the opposite side of the active site. Additionally, the % residues engaged in α-helices are also higher and more consistent in ShufPTP. This difference is primarily attributed to the increased length and stability of N-terminal helix. Despite this difference again appearing visually negligible, this extension is strongly linked to the interactions capable of restraining the base of the IPD-loop (further discussed in “IPD-Loop and P-Loop Dynamics in ShufPTP and TkPTP” section), therefore altering the dynamic behavior of the enzyme.

### Comparison of IPD-Loop and P-Loop Dynamics in ShufPTP and TkPTP

#### IPD-Loop Dynamics

Class I PTPs are characterized by the motion of the flexible loops decorating their active sites, with a mobile WPD-loop, that undergoes significant conformational transitions during catalysis (**Figure 1**), and a rigid phosphate binding P-loop.^91^ In contrast to the substantive acid loop motion observed in these PTPs, the analogous acid loop (the IPD-loop) in archaeal PTPs, including ShufPTP, is far rigid, preferring a closed or semi-closed conformation even in the PTP’s unliganded form (based on available structural information, see PDB IDs: 5Z59,^56^ 5Z5A^56^ and 7MPD^55^ as examples), while the P-loop instead is conformationally flexible^55, 56^ (**Figure 2**). Particularly, ShufPTP and TkPTP both demonstrate alternative conformations of their P-loops; however, despite their high sequence similarity, they demonstrate noticeable differences in their thermal stability and catalytic activities. We thus also compare the conformational dynamics of key catalytic loops in wild-type ShufPTP and TkPTP at both 300K and 360K, to understand both baseline dynamics and how they are affected by temperature.

**Figures 11** and **S9** show comparisons of IPD- and P-loop dynamics and conformational distributions in ShufPTP and in TkPTP, starting from the unphosphorylated enzyme and the phosphoenzyme intermediate, respectively, with the IPD-loop in a closed conformation at the start of the simulation. In the case of the unliganded simulations, we observe that the TkPTP IPD-loop is conformationally flexible (similar to dynamics observed in PTP1B in our prior simulations^31^). In contrast, and in agreement with our crystal structures of ShufPTP (in which the IPD-loop is primarily in a closed conformation, PDB IDs: 9E9N, 9E9L, 9E9M, 9E9U), the conformational ensemble of the IPD-loop (and even the P-loop), is less diverse and more restricted, and mostly concentrated on sampling catalytically optimal conformations associated with positioning D63 closer to the active site, while also stabilizing C93.

**Figure 11.**
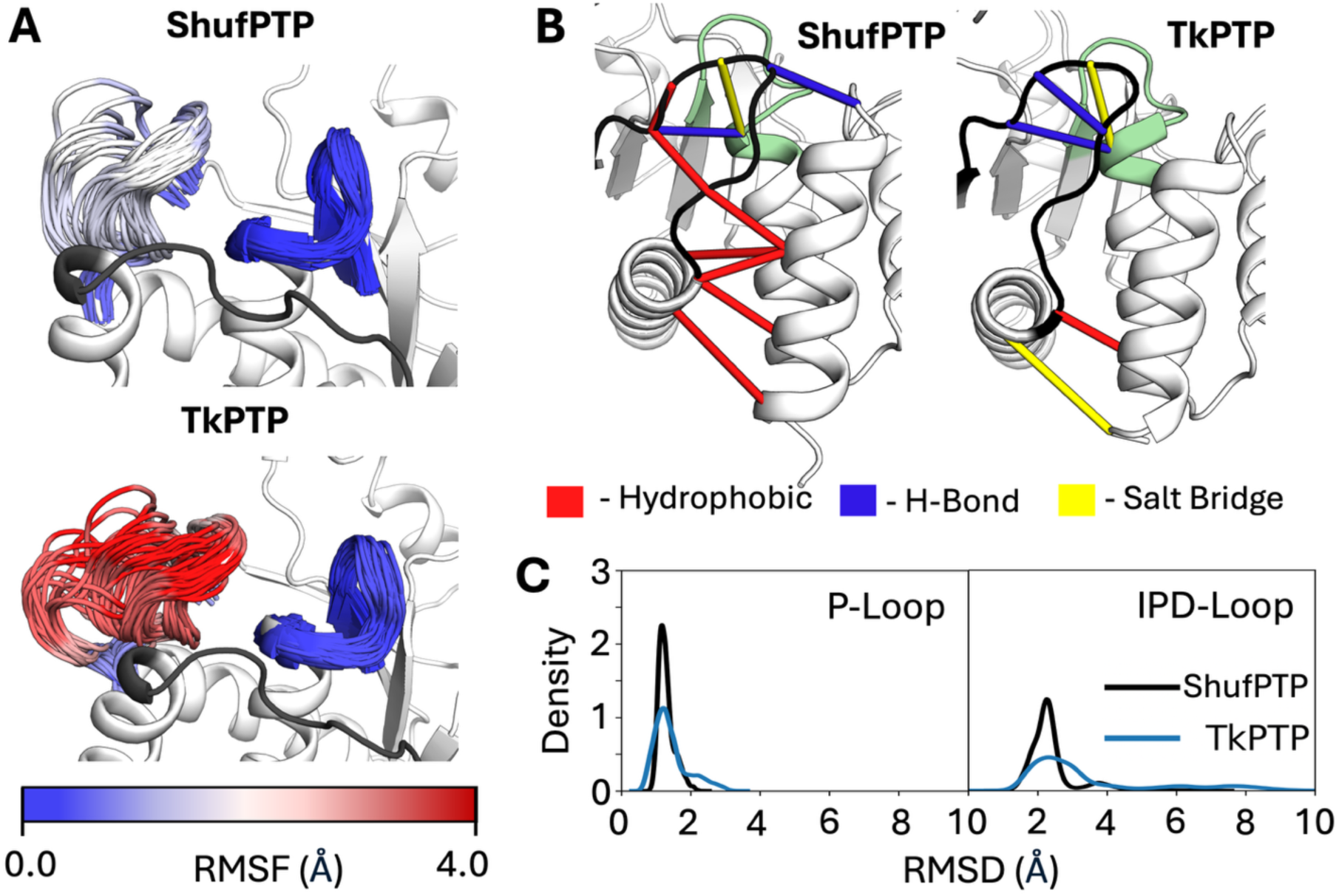
Conformational dynamics of the IPD- and P-loops in the unliganded forms of ShufPTP and TkPTP. Simulations were initiated from the IPD-loop-closed and low P-loop state of each enzyme. (**A**) The ensemble of sampled conformations during simulations of ShufPTP and TkPTP at the IPD-loop-closed unliganded state was visualized and colored based on the root mean square fluctuations (RMSF, Å) of loop C_α_-atoms. (**B**) A stabilizing hydrophobic network at the base of the IPD-loop was identified and compared to the interactions present in TkPTP. This network was obtained by calculations using Key Interaction Networks (KIN).^154^ (**C**) Assessment of the P-loop and IPD-loop conformational ensembles based on kernel density estimation (KDE) analysis of the root mean square deviations (RMSD, Å) of the backbone atoms of the IPD- and P-loops of ShufPTP and TkPTP. These data indicate a less conformationally diverse ensemble in TkPTP than in ShufPTP.

The restricted loop dynamics in ShufPTP can be partially rationalized by the extended C-terminal helix of the IPD-loop of ShufPTP, which carries additional hydrophobic residues such as F146, which is not present in TkPTP. This, in combination with a bulkier leucine at position 69 (rather than valine in TkPTP) reinforces a network of hydrophobic interactions that likely limit the opening of the IPD-loop and fix the conformation of this loop in its catalytically active closed conformation (**Figure 11B**). Further, our simulations at the phosphoenzyme intermediate state indicate that the P- and IPD-loop are rigidified, in what is likely a ligand-gated conformational change, as observed in our prior work on PTP1B and YopH.^31^ **Figure S9** shows that, in agreement with prior experimental work on TkPTP,^56^ the conformational dynamics of the P- and IPD-loops of TkPTP and ShufPTP exhibit temperature sensitivity. While both unliganded systems show broader conformational distributions at 360K than 300K, at the phosphoenzyme intermediate state, TkPTP exhibits a narrowing of its loop conformational ensemble with increasing temperature, while the corresponding conformational dynamics of ShufPTP remains largely unaffected.

#### P-Loop Dynamics

We further observe that, in agreement with experimental observations (ref. ^56^ and **Figure 2**), both TkPTP^56^ and ShufPTP show unusual conformational flexibility in their phosphate binding P-loops, which is particularly curious given that phosphate-binding P-loops tend to be rigid^31, 51-55, 155^ and fall within clearly defined ranges of conformational parameters (see *e.g.*, ref. ^155^ for characterization of the analogous Walker A P-loop). In particular, X-ray crystallographic structures of both TkPTP and ShufPTP show both catalytically active and inactive conformations of the phosphate-binding P-loops (**Figure 12C**), that differ by 0.4Å RMSD between their backbone atoms. In order to characterize transitions between these states in our MD simulations, the following root mean square deviation (RMSD)-like metric was used to compute the similarity between simulation frames extracted from our MD simulations, and the corresponding active/inactive conformations in the X-ray crystallographic structure (CS):

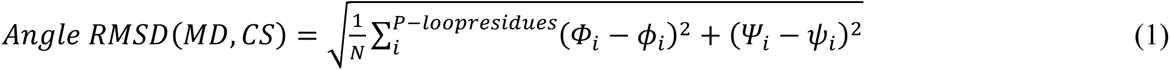

**Figure 12.**
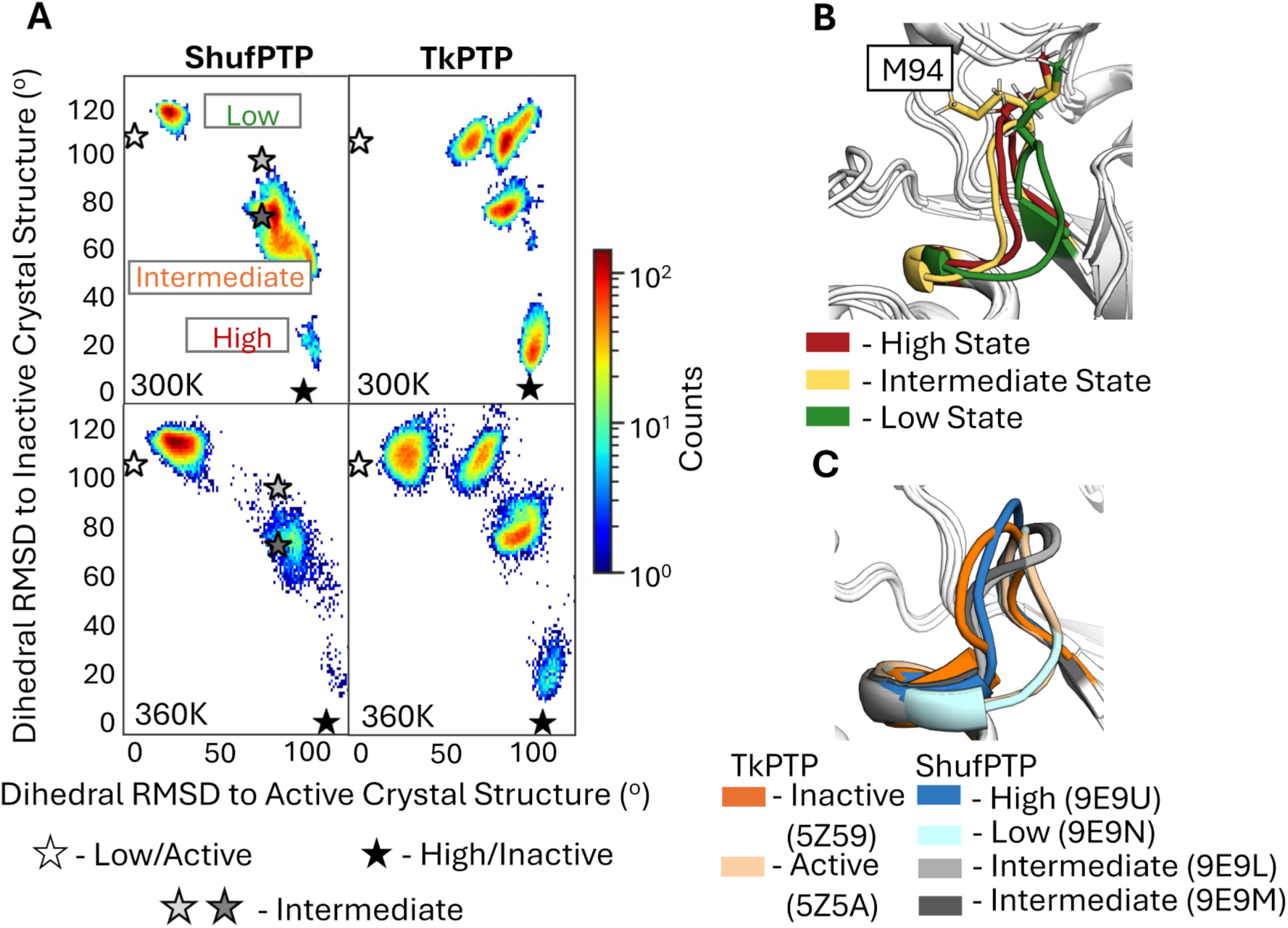
Comparison of P-loop conformational transitions between ShufPTP and TkPTP. (**A**) 2D histograms of the P-loop conformation sampled in our MD simulations of each enzyme, defined based on a root mean square deviation (RMSD)-like metric (Eq. 1 of the main text). The deviation between active (low) and inactive (high) P-loop state X-ray crystal structures (CS) of ShufPTP and TkPTP corresponds to φ and ψ angles of 116.1 degrees and 104.7 degrees respectively. The angle RMSD of two ShufPTP intermediates to active and inactive CS are: 9E9L = (78.6, 95.2) degrees, and 9E9M = (78.6, 71.6) degrees. Simulations were initiated from inactive (high) P-loop states of each enzyme, with the IPD-loop in its loop-closed state. Corresponding analysis initiated from the intermediates and active (low) P-loop state is shown in **Figures S10** and **S11**. The corresponding crystallographic positions of the P-loop are denoted by stars, as annotated on the figure. (**B**) Illustration of structures from the local maxima of the three states observed on the histogram of ShufPTP, in order to illustrate the key structural differences between the MD intermediate and the inactive (high) state highlighted. (**C**) Comparison of active (low), intermediate and inactive (high) P-loop conformations of TkPTP and ShufPTP, based on crystallographic data (PDB IDs: 5Z5A,^56^ 5Z59,^56^ 9E9N, 9E9L, 9E9M and 9E9U).

Here, *N* is the total number of P-loop residues considered, (𝛷*_i_,* 𝛹*_i_*) are backbone angles of the residue *i* in the reference crystal structure and (𝜙*_i_*, 𝜓*_i_*) are corresponding angles from the simulation frame. This analysis was then applied to simulations of wild-type ShufPTP and TkPTP initiated from active (low), intermediate and inactive (high) conformations of the P-loop, with the IPD-loop in its loop-closed conformation, in agreement with existing structural data (ShufPTP PDB IDs: 9E9N, 9E9L, 9E9M and 9E9U, this work, and TkPTP PDB IDs: 5Z59 and 5Z5A^56^).

When simulations are initialized from the active (low) P-loop state, both ShufPTP and TkPTP sample highly stable localized conformations of the P-loop, which aligns with the catalytic importance of this conformation (**Figure S10**). **Figure 12** illustrates that, in comparison to TkPTP, the inactive (high) state of the ShufPTP P-loop is severely destabilized and demonstrates a full transition towards an active (low) state at 300K. Even in the 360K simulations ShufPTP samples fewer intermediate conformations and demonstrates a rapid transition towards an active state. Specifically, the ShufPTP P-loop rapidly adopts an intermediate conformation between the two states, where the M94 side chain is rotated towards the active site of this PTP, further driving the transition towards the low P-loop state (**Figure 12B**). This transitional state observed in MD is in good agreement with crystallized intermediates, particularly the intermediate without vanadate at the active site (PDB ID: 9E9M) (**Figure S10**). In contrast, TkPTP explores a variety of intermediate conformations during our simulations, but the active conformation is reached only in the conditions of elevated temperature. The fact that we observe these transitions in ShufPTP in the 300K simulations but not in TkPTP already on the 500ns MD simulation timescales suggests a lower thermodynamic barrier between the inactive and active conformation of the ShufPTP P-loop, which would (1) partially explain its increased catalytic activity at room temperature (**Table 2**), and (2) explain why the TkPTP P-loop transition appears to be temperature dependent,^56^ in order to surmount this barrier. These differences are particularly intriguing, given the fact that ShufPTP and TkPTP have identical P-loop sequences (**Figure 3**).

Finally, simulations of ShufPTP with C93 in either its S-hydroxycysteine or its protonated forms showed stabilization of the opposite, high or inactive state *via* the crystalized intermediates (**Figure S12**). This is achieved through the flexibility of the P-loop, which allows for an extended position of R99 that stabilizes the closed state of the IPD-loop in the absence of phosphate. These observations place special importance onto the intermediate P-loop state, which, as was discussed in the section “Exploring Cysteine Oxidation in PTPs”, is also the state most susceptible to the solvation and further oxidation. This illustrates the potential of cysteine protonation changes to drive active site dynamics, exposing oxidation-prone intermediate P-loop conformations.

## Overview and Conclusions

Protein tyrosine phosphatases have proven to be an excellent model system for understanding the links between loop dynamics, catalysis, and enzyme evolution. These enzymes are regulated by the motion of a catalytic acid loop, that carries an evolutionarily conserved aspartic acid important for acid-base catalysis in PTPs.^91^ Structural, NMR, and molecular simulation studies have emphasized the links between loop motion and catalysis in these enzymes.^31, 47, 50, 91, 156, 157^ The more broad role of loop dynamics in enzyme evolution and protein design has also been a topic of increasing interest (see discussion in refs. ^35, 36^ and references cited therein, as well as other examples of more recent studies, *e.g.*, refs. ^34, 158-162^).

While PTPs as a family have been studied extensively,^91, 163, 164^ archaeal PTPs have been much less focused on, and of those that have been studied, trends demonstrate unusual biochemical and biophysical properties, including hyper-thermostability, catalytic backups, and mobile phosphate binding loops.^55, 56, 81^ A better understanding of archaeal PTPs thus plays a dual role both in expanding our understanding of loop dynamics and catalysis in PTPs and other enzymes regulated by catalytic loop motion more broadly, but also of how enzymes evolve to operate under inhospitable conditions, given that archaea are extremophiles.^165^ In this work, we have generated an extremophile chimeric PTP, based on amino acid shuffling of five hyper-thermophilic archaeal PTP sequences (**Figure 3**), which we denote ShufPTP, and extensively biochemically, biophysically, structurally, and computationally characterized this enzyme.

Archaeal enzymes are known to have unique structure-function properties,^166-168^ and ShufPTP is no exception. Dynamically and in terms of stability, calorimetry indicates that ShufPTP is extremely thermostable, with denaturation likely occurring at temperatures > 130 °C (**Figure 6**). This is particularly noteworthy, given that the highest recorded temperature for the growth of an archaeal strain, from *Methanopyrus kandleri*, is 122 °C.^165^ For comparison, previously characterized archaeal PTPs such as TkPTP and SsoPTP have *T*_m_ and activity at temperatures 65 °C or above,^55, 56^ and the source organisms for the PTPs used for the amino acid shuffling (**Figure S1**) have growth ranges between 50 – 95°C, and optimal growth temperatures in the 80 – 88 °C range.^84-88^ Although impressive, these temperatures are still substantively lower than > 130 °C for ShufPTP. Sequence comparison of thermophilic and mesophilic PTPs shows a general trend of higher frequency of Arg, Glu and Leu residues, lower frequency of Cys and Ser residues, higher hydrophobic content, and higher instability, aliphatic and GRAVY indexes, consistent with prior analysis, consistent with expectations,^144-147, 149, 150^ (**Supplementary Data**, the exception to this being the atypical PTP VHZ which has properties of both human and archaeal PTPs). However, our sequence analysis only allowed us to differentiate between thermophilic and mesophilic PTPs in broad terms. Detailed analysis of MD simulations of ShufPTP, TkPTP, YopH and PTP1B indicated that ShufPTP additionally maintains a higher % of residues engaged in hydrophobic contacts, as well as having a higher mean content of residues in α-helical structures over the course of our simulations (**Figure 10**), providing a further physical and structural basis for the increased thermostability of ShufPTP.

Following from this, and similarly to structural data available on other archaeal PTPs,^55, 56^ ShufPTP has a rather rigid acid loop (**Figures 4** and **11**), in contrast to the mobile acid loops observed in other PTPs.^91^ In contrast, the phosphate binding loop of ShufPTP is highly mobile, taking on at least three distinct stable conformations (**Figures 4** and **12**). This is a feature that is highly uncommon in P-loop containing enzymes more broadly, given that phosphate binding P-loops tend to be structurally rigid and follow narrowly defined geometric parameters.^31, 51-55, 155^ We note, however, that both TkPTP and SsoPTP have been observed to have mobile P-loops,^55, 56^ although to a lesser extent than we present here in the case of ShufPTP (**Figures 4** and **11**).

Mechanistically, the active site cysteine of ShufPTP is readily amenable to oxidation (**Figure 4**), a feature that has emerged to be an important regulatory mechanism that links cellular tyrosine phosphorylation with signaling by reactive oxygen or nitrogen species.^74^ Compared to other PTPs such as PTP1B, YopH and TkPTP, we observe greater solvation of the nucleophilic cysteine of ShufPTP in our simulations, likely leading to the more facile oxidation of this cysteine in ShufPTP (**Table 2**). Further, the oxidation of this side chain appears to, in turn, increase the mobility of the phosphate binding P-loop of ShufPTP (**Figure S12)**. The propensity of PTP active site cysteines towards oxidation has been linked to reactive oxygen species-mediated signal transmission.^169^ This can be further linked to the adaptation of this catalytic machinery to perform redox chemistry in nature, specifically in arsenate reductases that reduce arsenate to arsenite (see *e.g.*, refs. ^170-181^ for examples), and is unlikely to be a “fluke” in ShufPTP.

ShufPTP is also catalytically versatile, utilizing a backup mechanism involving a glutamic acid on a mobile loop as a redundancy for its preferred Asp-as-base mechanism (utilizing the aspartic acid on the acid loop, **Figures 7** and **8**). The presence of catalytic backups appears to be a common feature of archaeal PTPs, having previously been suggested for both TkPTP (E132)^56, 81^ and SsoPTP (E40),^55^ as well as, curiously, in the human dual specificity phosphatase VHZ (E134).^82^ In TkPTP and VHZ, the alternate catalytic mechanism is at least supported by the mutation of a typically invariant glutamine on the Q-loop that is essential for nucleophile positioning in the second step of the catalytic mechanism^134^ to a glutamic acid, whereas SsoPTP retains the glutamic acid on the Q-loop and uses an alternative residue, E40, on a nearby mobile loop as a backup. While the presence of catalytic backups is highly atypical for PTPs more broadly, and this is only the fourth documented example of such, backup mechanisms have been seen before in other enzymes. These include an organophosphate hydrolase, serum paraoxonase 1 (PON1),^135, 136^ as well as quorum-quenching (*QQ*) lactonases.^137^ PON1 and *QQ* lactonases are, however, scavenger enzymes that are functionally diverse, and thus it stands to reason that they would not only be substrate/catalytically promiscuous but also mechanistically promiscuous. Here, we show a further example of mechanistic promiscuity in an enzyme from a family of enzymes that is involved in the regulation of core cellular processes,^91^ where fine-tuning activity and selectivity is crucial. We demonstrate through simulations that tight regulation of the relative mobility of the loops decorating the active site facilitates interchanging catalytic mechanisms.

Each of these features alone would render ShufPTP a highly unusual enzyme. Taken together, it is a perfect illustration of the extreme properties observed in archaeal enzymes more broadly.^166-168^ Despite this high similarity to extant PTPs, ShufPTP shows much more extreme biochemical and biophysical properties than either of the previously characterized archaeal PTPs, TkPTP^56^ and SsoPTP,^55^ to which ShufPTP shares 85 and 39% sequence identity, respectively. The fact that archaeal proteins, which have adapted to extreme conditions of temperature, pressure and salinity, can have unusual biophysical properties has been well documented.^166-168, 182^ Our data showcase how a simple jump in evolutionary space can radically alter an archaeal enzyme while maintaining high levels of the native activity, and simultaneously greatly diversifying the enzyme’s biophysical profile. This further underscores the potential of engineered archaeal enzymes in biotechnology, which is an area of rapidly growing interest.^166, 167, 183-187^

## Materials and Methods

### Chemicals

Dithiothreitol (DTT) and ampicillin (AMP) were purchased from GoldBio. Protease-inhibitor tablets were purchased from Sigma-Aldrich. All other buffers and reagents were purchased from Sigma-Aldrich or Fisher. The substrate p-nitrophenyl phosphate (*p*NPP) was synthesized using published methods.^188^ Crystallography screens, trays, and coverslips were purchased from Hampton Research.

### Protein Expression and Purification

The plasmid pET-21+ encoding ShufPTP and its variant D63N were synthesized by Twist Bioscience (San Francisco, CA). The DNA was transformed into *Escherichia coli* BL21-DE3 cells and grown overnight at 37 °C on Luria-Bertani (LB) culture plates containing 100 µg mL^−1^ ampicillin. One colony was selected and placed into 10 mL of SOC media containing 100 µg mL^−1^ ampicillin and grown overnight. The following morning, 1 L of LB media containing 100 µg mL^−1^ ampicillin was inoculated with the 10 mL of overnight growth and shaken at 170 rpm at 37 °C until the optical density at 600 nm (OD600) reached 0.6–0.8. After the optimal OD was reached, the 1 L growth was induced by 0.1 mM isopropyl β-D-thiogalactoside (IPTG) and shaken at 170 rpm at room temperature overnight. The cells were harvested by centrifugation at 12,000g for 30 min at 4 °C and stored at −80 °C.

ShufPTP cells were thawed on ice and resuspended in 10x their equivalent volume of lysis buffer (50 mM imidazole pH 7.5, 1 mM ethylenediaminetetraacetic acid (EDTA), 7 mM dithiothreitol (DTT), 5 mM tris(2-carboxyethyl)phosphine (TCEP), and 10% glycerol) supplemented with protease inhibitors (0.5 mg mL^−1^ aprotinin, 0.7 mg mL^−1^ pepstatin, and 0.5 mg mL^−1^ leupeptin). The cells were lysed by sonication. The cell lysate was centrifuged at 29,000g for 30 min at 4 °C. The supernatant was heated to 70 °C for 15 minutes and filtered with a 0.45 μm syringe filter.

The filtrate was purified using a 5 mL HiTrap Q HP column equilibrated with lysis buffer on a fast protein liquid chromatography (FPLC) system. The cell lysate was loaded at 1.5 mL min^−1^ and washed with lysis buffer until the absorbance at 280 nm reached baselined. Flow-through fractions exhibiting absorbance at 280 nm were collected and tested with para-nitrophenyl phosphate (pNPP) for phosphatase activity. Active fractions were analyzed for purity using SDS-PAGE on 20% gels.

The active fractions were pooled (30-40 mL), concentrated to less than 12 mL, and loaded onto a pre-equilibrated HiLoad 26/60 Superdex 200 prepgrade column (GE) equilibrated with a buffer containing 30 mM Tris buffer pH 7.5 containing 25 mM sodium chloride, 0.2 mM EDTA, 7 mM DTT, and 5 mM TCEP. Fractions were assayed with pNPP for activity and analyzed for purity using 20% SDS-PAGE. Pure protein was concentrated to 5–10 mg mL^−1^ and either used immediately for crystallization or supplemented with 10% glycerol, flash-frozen in liquid nitrogen, and stored at −80 °C in aliquots.

### Crystallization and Structure Determination

Crystals for ShufPTP were grown by hanging drop vapor diffusion using 8 mg mL^−1^ protein and a precipitant solution of 0.1 M CAPS pH 10.5, and 30% PEG 400 at a 1:1:0.2 protein : well : 20% additive screen drop ratio. The additive screen contained 0.2 M ammonium sulfate ((NH_4_)_2_SO_4_), 20-30% polyethylene glycol 4000 (PEG 4000), and 10-35% glycerol. The vanadate bound structure was obtained by adding 5 mM sodium metavanadate (Na_3_VO_4_) to the protein for co-crystallization. Crystals were transferred to a cryo-protectant solution containing mother liquor, 20% additive screen, and 10-15% glycerol before flash freezing in liquid nitrogen.

Diffraction data were collected on the Stanford Synchrotron Radiation Lightsource (SSRL) beamline 9-2 or on a home source. The total radiation dose on the crystals collected at SSRL is estimated to be below 25 MGy as calculated by RADDOSE-3D.^189^ Data were indexed and processed using DENZO and SCALEPACK in the HKL3000 program suite.^190^ Molecular replacement was performed with Phaser-MR as implemented in Phenix using wild-type TkPTP (PDB ID: 5Z5A) as a search model. Phenix.refine (version 1.21.2-5419)^191^ was used for refinement. Model building was performed using Coot.^192^ All figures of the enzyme structures and structural alignments therein were made using PyMOL.^193^ Representative 2fo-fc electron density maps for each structure are shown in **Figure S13**. Simulated annealing omit maps for the oxidized cysteine residues are shown in **Figure S2**. Data collection and refinement statistics are reported in **Table S1**.

### Differential Scanning Calorimetry (DSC)

DSC experiments were carried out using a VP-DSC calorimeter from MicroCal, following protocols we have described previously in detail.^194^ Briefly, protein solutions for the calorimetric experiments were prepared at a concentration of 1 mg/mL in 50 mM sodium acetate, 100 mM Tris, 100 mM Bis-Tris pH 7. The same buffer was used to fill the reference cell of the calorimeter. All DSC scans were carried out under an overpressure of about 2 atm and at a scanning rate of 1.5 degrees per minute. Before each experiment with protein solution in the sample cell, several buffer-buffer baselines were recorded to ensure calorimeter equilibration.

### Steady-State Kinetics

Steady-state kinetic parameters were measured at temperatures from 22 to 90 °C. For the pH-rate profiles, concentrated aliquots of ShufPTP and its variant D63N were thawed on ice and diluted with a buffer base mix (BBM) containing 50 mM sodium acetate, 100 mM Tris, and 100 mM bis-Tris from pH 4.0 to pH 6.25. This buffer system maintains constant ionic strength throughout the pH range examined. For the Arrhenius plot, aliquots of ShufPTP were diluted with the same buffer mix at the pH optimum of 4.75. At this pH the primary buffering component is acetate, which shows a minimal temperature effect on its p*K*_a_. All kinetic measurements were performed in the presence of 2 mM DTT, except for the cysteine oxidation measurements described below.

A 50 mM solution of the disodium salt of *p*NPP was prepared in the same buffer mix. The reactions were run on 96-well plates using diluted enzyme and a range of substrate concentrations. Reactions were allowed to proceed for 5 min and quenched using 50 μL of 5 M NaOH, and the amount of the product *p*-nitrophenol was assayed from the absorption at 400 nm using the molar extinction coefficient of 18,300 M^−1^ cm^−1^. Reaction blanks were made using identical conditions, replacing the enzyme with buffer to correct for non-enzymatic hydrolysis of the substrate. The amount of product released and elapsed time were used to calculate the initial rates. The data were fitted to the Michaelis–Menten equation to obtain the kinetic parameters. Kinetic data were obtained on both ShufPTP and its variant D63N as a function of pH to obtain pH-rate profiles. The bell-shaped pH-rate profiles were fitted to Eq 1, the standard equation relating the dependence of the observed *k*_cat_ to the maximal, or limiting, value as a function of pH, where catalysis is dependent on two ionizable residues, one protonated and the other deprotonated.^195^

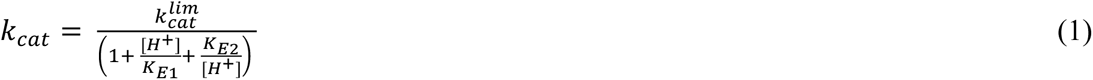

### Active-Site Cysteine Inactivation using Hydrogen Peroxide

Enzyme stock was diluted in BBM at pH 4.75 with 0.1-0.8 mM hydrogen peroxide. Reaction blanks were made with the same reagents omitting the hydrogen peroxide. The enzyme was incubated with peroxide, and aliquots were removed to assay for activity with 7.6 mM *p*NPP. Phosphatase activity assays proceeded for 5 minutes in BBM at pH 4.75 and quenched with 5 M NaOH. The formation of *p*-nitrophenol product was assayed as described above.

At each hydrogen peroxide concentration, residual activities as a function of time were plotted to obtain *k*_obs_ (s^−1^) for the inactivation rate constant at hydrogen peroxide concentration. These *k*_obs_ values were plotted against hydrogen peroxide concentrations to obtain the second-order rate constant *k*_inact_ (M^−1^ s^−1^). The same methodology was used to obtain the *k*_inact_ for YopH and PTP1B.^98^

### Substrate Preference Determination using an NMR-based competitive method

These experiments and NMR analyses were conducted following our recently described methodology.^115^ A substrate mixture containing 5 mM of both α- and β-naphthyl phosphate were made in BBM pH 5.0. In separate experiments at 22 and 60 C, ShufPTP at a concentration of 0.7 µM was allowed to react with the substrate mixture until the fraction of reaction reached approximately 20%. The reactions were then quenched with 50 μL 1 M NaOH, raising the pH beyond the range of enzymatic activity. Phosphate esters are extremely unreactive, and control experiments showed no detectable spontaneous hydrolysis. 500 μL of each quenched reaction mixture, at room temperature, was added to an NMR tube with a stem coaxial insert tube containing phenyl phosphonic acid at a concentration of 50 mM in D_2_O. The operational frequency for ^31^P collection was 202.46 MHz and ran at 296.2 K. ^31^P 90° pulse widths were measured using a pulse delay of 60 seconds and inverse gated decoupling (Bruker pulse sequence: zgig) to avoid NOE effects. ^31^P T1 measurements were performed using the optimized 90° pulse and inverse gated decoupling (Bruker pulse sequence: t1irig). Chemical shifts were calibrated by external standard (H_3_PO_4_) using a co-axial tube to ensure buffer integrity.

Kinetic mixture samples were run with 32 scans, a relaxation delay of 35 seconds, an offset frequency of 6ppm, a spectral width of 46 ppm, and a 32k points collection. Processing was performed on the software package MestReNova 14.2. The data was zero-filled to 64k points, and a line broadening factor of 1 Hz was applied. Peak areas were measured using quantitative global spectra deconvolution with 4 fitting cycles.

### Molecular Dynamics Simulations

Molecular dynamics (MD) simulations were performed to simulate the dynamical behavior of TkPTP, ShufPTP, and the ShufPTP D63N variant, considering both the active (PDB ID: 5Z5A^56^ and 9E9U) and inactive (PDB ID: 5Z59^56^ and 9E9N) forms of the P-loop. Additionally, ShufPTP was simulated in its unphosphorylated intermediate forms (PDB IDs: 9E9L and 9E9M), active form with C93 in its S-hydroxycysteine and protonated states, and inactive form with C93 in its S-hydroxycysteine and protonated states, to obtain a comprehensive description of P-loop dynamics. The IPD-loop is in its closed conformation in all available crystal structures. Simulations of TkPTP and ShufPTP were initiated from their respective crystal structures; in the absence of crystallographic data, D63N ShufPTP was constructed using the PyMOL^193^ Mutagenesis Wizard (selecting the Asn rotamer most similarly located to the catalytic Asp in the TkPTP structure).

Simulations were performed both in the unliganded and phosphoenzyme intermediate states of structures with an active (low) P-loop conformation, and in only the unliganded state of structures with inactive (high) and intermediate conformation of their P-loop (as this conformation does not accommodate the phosphoenzyme intermediate). We note that the cysteine in the ShufPTP crystal structures is in an oxidized form; to mimic a catalytically active state of this enzyme, the C93 side chain was converted back to its reduced form using the PyMOL^193^ Mutagenesis Wizard. As two rotamers of crystalized C93 were observed in the inactive and intermediate crystal structures of ShufPTP (**Figure 4**), the rotamer in the catalytically active conformation of this side chain was chosen to represent reduced cystine. In the case of the unliganded simulations, the catalytic cystine was deprotonated, and in the simulations of the phosphoenzyme intermediate, PO_4_ group was added manually. Both structural modification were handled *via* the CHARMM-GUI.^196^ Further, the oxidized cysteine in the ShufPTP crystal structure pushes out the D63 side chain such that it forms a catalytically non-productive dyad with the E41 side chain (**Figure 4**). To mitigate this issue, we again used the PyMOL^193^ Mutagenesis Wizard to select the D63 rotamer that was in the closest analog to the position of this side chain in TkPTP as the starting point for our simulations. All molecular dynamics (MD) simulations were performed using the GPU-accelerated GROMACS 2024.^197^ All simulations were conducted using CHARMM36m^198^ force field and TIP3P water model, at both 300 and 360K (as the enzymes of interest are thermophiles). In total, 18 different systems were simulated: 3 enzyme variants, in both unliganded and phosphoenzyme intermediate states, with the unliganded systems further simulated in P-loop active (low) and inactive (low) conformations (the phosphoenzyme intermediate was only simulated in the P-loop active “low” conformation). For each system, 5 independent trajectories of 500 ns each were propagated using different initial velocities assigned using different random seeds, using a 2 fs time step, leading to a total of 45 μs of cumulative simulation time.

All systems were prepared for simulation using the CHARMM-GUI,^196^ and followed a standard energy minimization, heating (NVT ensemble) and equilibration (NPT ensemble). Production simulations were all performed in the NPT ensemble (1 atm pressure). All simulation analysis was performed using MDAnalysis 2.7.0,^199, 200^ with the exception of non-covalent interaction analysis, which was performed using Key Interaction Networks.^154^

Solvent thermodynamics near the active site of ShufPTP and TkPTP were analyzed using Grid Inhomogeneous Solvation Theory (GIST).^120, 121, 201^ Low/active, intermediate and high/inactive starting structures were used to initialize 3x100ns simulations of each system. The protocol of restrained MD using Amber24 and GIST analysis was performed following the protocol of ref. ^121^. GIST analysis was performed using AmberTools24^202^ and the gisttools analysis package (https://github.com/liedllab/gisttools). The free energy of solvation of the protein (ΔA_solv_) was calculated for a 50Åx50Åx50Å grid centered on the S_γ_-atom of C93, from which ΔA_solv_ was integrated over voxels located within a 5Å sphere of the S_γ_-atom of C93.

Full details of simulation and analysis protocols are provided in the **Supporting Information**, and a data package containing sample input files, starting structures, any non-standard parameter files, representative simulation snapshots, and any custom simulation analysis scripts, is provided on Zenodo at the following DOI: 10.5281/zenodo.15074903.

### Empirical Valence Bond Calculations

The rate-limiting hydrolysis of the phosphoenzyme intermediates of ShufPTP and its variant D63N were described using the empirical valence bond (EVB) approach,^138^ following our prior work.^31, 33, 203^ Here, we have performed EVB simulations of the hydrolysis step of the reactions facilitated by either D63 or E132 (D63-as-base and E132-as-base), to compare the energetic barriers of the dominant and proposed backup mechanisms. These reactions were modelled using the valence bond states presented in Figure S19 of ref. ^31^. Note that as both reactions involve proton abstraction by a carboxylate side chain, identical EVB parameters and valence bond states were used to describe the two mechanistic possibilities. Further, we used the same EVB parameters as presented in prior work^204^ for the current calculations.

All EVB simulations were performed using the catalytically active (low) conformation of the ShufPTP P-loop (PDB ID: 5Z5A, 9E9N). The system preparation and initial equilibration for EVB simulations were performed as described in the **Supporting Information**. Each EVB trajectory was initialized from 20 independent snapshots extracted from the corresponding MD simulations of wild-type and D63N ShufPTP. Starting structures for each EVB trajectory, which are provided on Zenodo (DOI: 10.5281/zenodo.15074903), were selected based on a catalytic distance cutoff of 5Å between the terminal O atoms of D63/E132 and the P atom of the phosphorylated C93 side chain. Each trajectory was first equilibrated for 20 ns at the approximate EVB transition state (*λ* = 0.5), with the subsequent EVB trajectories propagated from the transition state in both the reactant and product directions, as described in our previous work.^31, 204^ Each EVB simulation was performed in 51 individual mapping windows of 200 ps in length per trajectory. This led to a total of 20 ns of equilibration and 10.2 ns EVB simulation time per trajectory, 400 ns equilibration and 204 ns EVB simulation time cumulative per system, and 2μs equilibration time and 1.02 μs EVB simulations over all 5 systems studied (four different starting states / 2 mechanisms for wild-type ShufPTP, and one starting state/mechanism for D63N ShufPTP, as summarized in **Table S3**).

All EVB simulations were performed using the *Q6* simulation package^205^ and the OPLS-AA^206^ force field, for consistency with previous related studies. All EVB parameters necessary to reproduce our work, as well as a detailed description of the computational methodology and subsequent simulation analysis can be found in the **Supporting Information**, and on Zenodo, DOI: 10.5281/zenodo.15074903.

## Supporting Information

Additional computational methodology, simulation details, and raw data for key figures. All parameters and files necessary to reproduce the simulations are available on Zenodo under DOI: 10.5281/zenodo.15074903.

## Acknowledgments

This material is based upon work supported by the National Science Foundation under Grant Nos. 2414074 and 2414075, by the Knut and Alice Wallenberg Foundation under Grant No. 2019.0431, and by the Swedish Research Council under Grant No. 2019-03499. JPL was supported by NIH GM112781. NRM and EAG were supported by NSF grant 2032315. The computations were enabled by resources provided by the National Academic Infrastructure for Supercomputing in Sweden (NAISS), partially funded by the Swedish Research Council through grant agreement no. 2022-06725. We acknowledge the National Academic Infrastructure for Supercomputing in Sweden (NAISS), partially funded by the Swedish Research Council through grant agreement no. 2022-06725, for awarding this project access to the LUMI supercomputer, owned by the EuroHPC Joint Undertaking and hosted by CSC (Finland) and the LUMI consortium. This work used the Anvil supercomputer at Purdue University through allocation BIO240204 from the <u>Advanced Cyberinfrastructure Coordination Ecosystem: Services & Support</u> (ACCESS) program,^207^ which is supported by U.S. National Science Foundation grants #2138259, #2138286, #2138307, #2137603, and #2138296. Further calculations were performed on the Theta and Polaris clusters at the Argonne Leadership Computing Facility, supported by a Director’s Discretionary award. Use of the Stanford Synchrotron Radiation Lightsource, SLAC National Accelerator Laboratory, is supported by the U.S. Department of Energy, Office of Science, Office of Basic Energy Sciences under Contract No. DE-AC02-76SF00515. The SSRL Structural Molecular Biology Program is supported by the DOE Office of Biological and Environmental Research, and by the National Institutes of Health, National Institute of General Medical Sciences (P30GM133894). The contents of this publication are solely the responsibility of the authors and do not necessarily represent the official views of NIGMS or NIH. Finally, the authors would like to thank Darko Mitrovic and Lucie Delemotte for helpful discussion and initial simulations.

