## Supplementary Information for "Conformational Dynamics and Catalytic Backups in a Hyper-Thermostable Engineered Archaeal Protein Tyrosine Phosphatase"

### Table of Contents

S1. Computational Methodology ........ S3

Molecular Dynamics Simulations ........ S3

Molecular Dynamics Analysis ........ S4

Grid Inhomogenous Solvation Theory (GIST) Analysis ........ S6

S2. Supplementary Figures ........ S10

S3. Supplementary Tables ........ S26

S4. Supplementary References ........ S29

### S1. Computational Methodology

##### Molecular Dynamics Simulations

Molecular dynamics (MD) simulations of TkPTP, ShufPTP, and the ShufPTP D63N variant, were performed using GPU-accelerated GROMACS 2024,[1] with system preparation performed as described in the main text. Enzymes with the active (low) P-loop form (PDB IDs: 5Z5A[2] and 9E9N) were simulated in the unliganded and phosphoenzyme intermediate states, while crystal structures with the inactive (high) (PDB IDs: 5Z59[2] and 9E9U) and intermediate (PDB IDs: 9E9L, 9E9M) P-loop were simulated in their unliganded state. Additionally, to further investigate P-loop dynamics of ShufPTP, active/low and inactive/high crystal structures were simulated with the C93 side chain in in its S-hydroxycysteine and protonated states. To account for temperature effects, simulations were performed at both 300 and 360K in the NPT ensemble (1atm), leading to 18 discrete simulation systems.

All simulations were performed using the CHARMM36m force field,[3] and the TIP3P water model,[4] using a common equilibration and production protocol described below. Each system was simulated for 5 x 500 ns of production using different initial velocities (assigned using different random seeds) per trajectory, leading to a total of 45 µs of cumulative simulation time over 18 systems. System preparation was performed using the standard protocol of the CHARMM-GUI[5] builder, with the phosphoenzyme intermediate parameterized using CHARMM-GUI solution builder tools. The parameters for the phosphoenzyme intermediate are presented in the associated Zenodo data package, DOI: 10.5281/zenodo.15074903. Each system was solvated in a cubic box of 67x67x67Å or 10Å from the edge boundary of the protein with 0.15M NaCl added to the system.

All systems were equilibrated using a common protocol, in the following steps: (1) 5000 steps of initial energy minimization using the steepest descent algorithm. (2) Gradual heating of the system from 100 to 300K (or 360K) over the course of 1.25 ns of simulation time in the NVT ensemble, during which backbone heavy atoms were restrained using 400 kJ $mol^{-1}$ $Å^{-2}$ harmonic restraints, while sidechain atoms were restrained using 40 kJ $mol^{-1}$ $Å^{-2}$ harmonic restraints. (3) A further 1 ns simulation in the NPT ensemble, in which the backbone heavy atom restraints were reduced to 100 kJ $mol^{-1}$ $Å^{-2}$, and the side chain restraints were removed. (4) The backbone restraints were dropped again to 50 kJ $mol^{-1}$ $Å^{-2}$ for another 1ns long NPT simulation. (5) A 1 ns long MD simulation without restraints was performed to complete the equilibration. Finally, 500 ns of production MD was performed per trajectory, in an NPT (1atm) ensemble. Simulation equilibration is shown in **Figures S14** to **S16**.

All NVT and NPT simulations used velocity rescaling[6] for temperature coupling, and all NPT simulations used a stochastic cell rescaling barostat[7] to control pressure. Both NVT and NPT equilibration simulations used the LINCS algorithm[8] to restrain all bonds to hydrogen atoms. All simulation steps used a 10Å non-bonded interaction cut-off to evaluate long-range electrostatic interactions, using the Verlet non-bonded cutoff scheme,[9] with long range electrostatics being evaluated using the Particle Mesh Ewald (PME) algorithm.[10] Finally, snapshots were saved every 100ps of simulation time for further analysis.

**Molecular Dynamics Analysis**

Analysis of loop dynamics and catalytic distances observed during our MD simulations was performed using MDAnalysis 2.7.0.[11, 12] Root mean square deviation (RMSD)[13] and fluctuation (RMSF)[14] analysis for all systems was conducted using backbone and $C_\alpha$-atoms, respectively. Further, RMSF values were visualized using simulation snapshots obtained every 5ns of simulation time, where the residue color is altered to represent a corresponding RMSF value *via* B-factor visualization in PyMOL 3.0.3.[15]

P-loop conformational analysis was performed by extracting the $(\phi, \psi)$ angles of each residue in the P-loop (residues 94-99) from every 100ps of the MD simulations initialized from the P-loop active/low (PDB: 5Z5A[2] for TkPTP, and 9E9N for ShufPTP), intermediate (PDB: 9E9L and 9E9M for ShufPTP) and inactive/high (PDB: 5Z59[2] for TkPTP, and 9E9U for ShufPTP) conformations. From this, an "angle RMSD" was calculated to assess the similarity between simulation frames extracted from the MD simulations and the corresponding crystallographic structure according to Equation 1, as described in the main text.

All distance analysis was performed by computing a minimum distance between the phosphorus atom of the phosphorylated C93 at the intermediate state, and the closest oxygen atom on the carboxylic acid side chain of the assessed residues (E38, E39, E41, D63, E132). Quantification of the presence of a potentially nucleophilic water molecule was performed by identifying the number of frames in which the O-P distance is less than 6Å, and a water molecule is located within 3.5Å of the carboxylic O and phosphate P atoms.

Non-covalent interaction analysis was conducted using Key Interaction Networks.[16] The interactions and their conservation scores in the active unphosphorylated simulations were calculated and compared between TkPTP and ShufPTP to identify uniquely stabilizing interactions. Solvent density, water content within the 5Å sphere around the catalytic cysteine and quantification of residues involved in α-helices were performed using MDAnalysis 2.7.0.[11, 12] A data package containing sample input files, starting structures, any non-standard parameter files, representative simulation snapshots, and any custom simulation analysis scripts is provided on Zenodo at the following DOI: 10.5281/zenodo.15074903.

**Grid Inhomogeneous Solvation Theory (GIST) Analysis**

Solvent thermodynamics within the active site of the low/active, intermediate and high/inactive P-loop states of ShufPTP and TkPTP were evaluated using GIST analysis.[17-19] Due to the absence of a structure of TkPTP in an intermediate P-loop state, and the requirement of strong solute restraints during the short MD simulations for the GIST analysis, high likelihood structures representative of the active, intermediate and inactive P-loop states were chosen from 5x500ns 300K unrestrained MD simulations of Tk-PTP and ShufPTP as starting points for the restrained simulations performed for the GIST analysis to ensure equivalent treatment of all starting structures.

All systems were simulated using TIP3P[4] water molecules and the Amber ff14SB[20] force field, as implemented in the Amber24 simulation package.[21] Solvated protein systems were energy-minimized with Cartesian positional restraints of 1.0 $kcal \cdot mol^{-1} \cdot Å^{-2}$ applied to the protein atoms and equilibrated in NVT ensemble. Equilibration was performed with Langevin dynamics at 300K and timestep of 1fs for 125ps. Bonds to hydrogen atoms were constrained using the SHAKE algorithm.[22] The 1.0 $kcal \cdot mol^{-1} \cdot Å^{-2}$ positional restraints were maintained on the protein throughout the simulations. Production simulations were run in NPT at 300 K and 1 atm using Langevin dynamics, Particle Mesh Ewald (PME) electrostatics,[10] and a 2 fs timestep with SHAKE[22] applied to bonds involving hydrogen atoms. Trajectories were 100 ns long per replicate (three replicates per state), in line with tutorial recommendations for obtaining converged GIST quantities.[18] Water configurations were saved every 2ps. All non-hydrogen protein atoms were harmonically restrained with positional restrains of 100.0 $kcal \cdot mol^{-1} \cdot Å^{-2}$.

All trajectories were centered on the $S_\gamma$-atom of C93 to ensure GIST grid formation around the active site. The GIST grid was constructed using 0.5 Å spacings and 100x100x100 voxels to capture bulk solvent located outside of the protein. Solvation free energy density of the active site was computed by calculating $\Delta A(r)$ based on the GIST outputs using Eq. S1:

$$\Delta A(r) = \Delta E_{WW}\,(\boldsymbol{r}) + \Delta E_{SW}\,(\boldsymbol{r}) - T\Delta S_{six}\,(\boldsymbol{r}) \tag{S1}$$

and integrating it over voxels that lie within a 5Å sphere around Cys93 S atom.

**Empirical Valence Bond Simulations**

*System Preparation for the Empirical Valence Bond Simulations*

Empirical valence bond (EVB) simulations[23] were performed to characterize the second, rate-limiting hydrolysis step of the reaction to analyze the relative catalytic roles of D63 located on the Acid loop and E132 located on the Q-loop of ShufPTP. This analysis was carried out for ShufPTP and its variant D63N, with a closed IPD-loop, and the P-loop in the active (low) conformation (PDB: 9E9N). To account for the dynamic nature of the electronic environment at the active site of the enzyme, we assessed the five following scenarios (**Figure S7**): the reaction in ShufPTP is catalyzed by D63 while E132 is located within or outside of the catalytic distance cutoff, the reaction is catalyzed by E132 while D63 is located within or outside of the catalytic distance cutoff, and the reaction in variant D63N, where E132 is the only remaining reactive residue (due to elimination of D63). All systems were simulated with 20 replicas, initialized from the frames of the MD simulations of phosphorylated ShufPTP with active (low) P-loop. Frames were selected using MDAnalysis, while setting the catalytic distance cutoff to 5Å. The reactive water was manually adjusted in each selected frame to ensure a catalytically productive conformation. Simulation starting structures can be found in the associated Zenodo package, DOI: 10.5281/zenodo.15074903.

All systems were solvated in a 23 Å radius droplet of TIP3P[4] water molecules, centered on the P atom of phosphorylated C93 of the P-loop, and described using the Surface Constrained All

Atom Solvent (SCAAS) model,[24] as in our previous work.[25-27] In this model, all residues within the inner 85% of the sphere are fully mobile during the simulations, all residues within the outer 15% of the sphere are restrained to their original crystallographic positions using 10 kcal $mol^{-1}$ $Å^{-2}$ harmonic positional restraints, and all residues outside the water droplet are constrained to their crystallographic coordinates using 200 kcal $mol^{-1}$ $Å^{-2}$ harmonic positional restraints. Residues outside the water droplet and in the restrained region of the droplet were simulated in their uncharged forms, in order to avoid introducing system instabilities associated with the presence of charged residues outside the explicit simulation sphere. A complete list of the assigned protonation states of all ionizable residues in our EVB calculations, as well as the histidine tautomerization states assigned, are provided in **Table S4**. All EVB parameters used in this study follow our prior work,[28] and have been provided in the associated Zenodo data package, DOI: 10.5281/zenodo.15074903.

*System Equilibration and Empirical Valence Bond Simulations*

All EVB simulations in this work were performed using the *Q6*[29] simulation package, using the OPLS-AA[30] force field as implemented into Q6. A 10 Å cut-off was set for all non-bonded interactions, except those involved in the chemical reactions, for which the cut-off was set to 99 Å (effectively no cut-off). Long-range electrostatic interactions were described using the local reaction field (LRF)[31] approach, and the temperature was controlled by the Berendsen thermostat.[32] All systems underwent a sequence of short equilibration and heating simulations over a total of 213 ps of simulation time, in order to gradually remove possible steric clashes and bad contacts in the system prior to equilibration at 300 K. Once the system reached 300 K, all restraints on the mobile region of the protein were removed, and only weak (0.5 kcal $mol^{-1}$ $Å^{-2}$) harmonic positional restraints were retained on the reacting moieties. Finally, we performed a further 20 ns of equilibration. Finally, weak harmonic positional restraints of 0.5 kcal mol$^{-1}$ Å$^{-2}$ placed on the reactive atoms during the equilibration were retained for the production EVB simulations as well. All further analysis of the data was done using *Q*tools v0.5.10 (Zenodo DOI: 10.5281/zenodo.842003).

### S2. Supplementary Figures

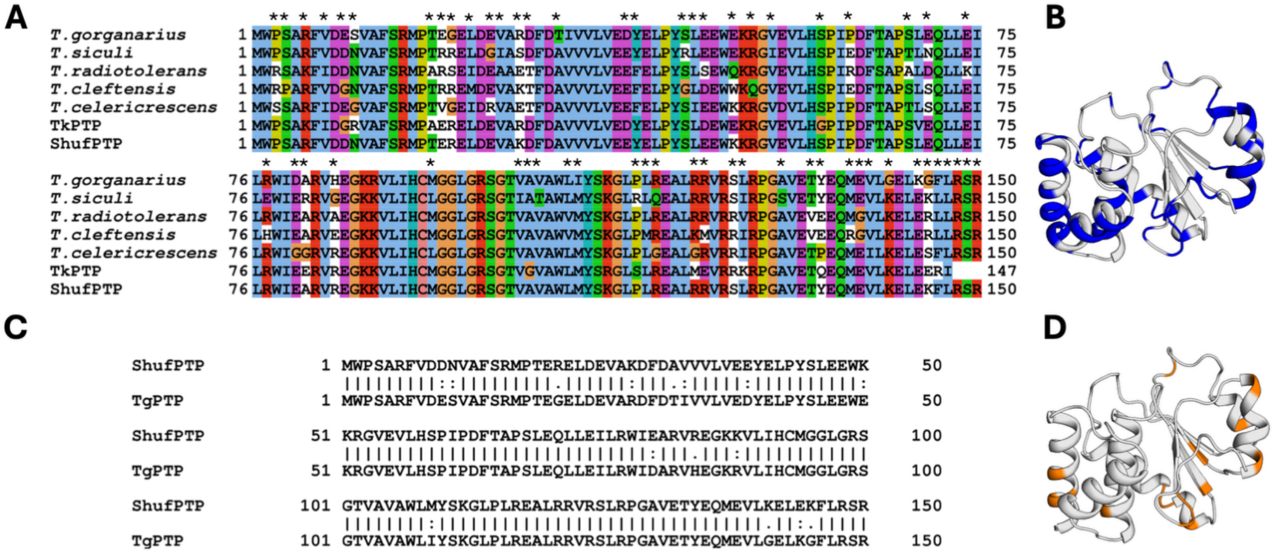


| | *T. radiotolerans* | *T. siculi* | *T. celericrescens* | *T. cleftensis* | *TkPTP* | *T. gorgonarius* | *ShufPTP* |
|---|---|---|---|---|---|---|---|
| ***T. radiotolerans*** | 100.0 | | | | | | |
| ***T. siculi*** | 76.0 | 100.0 | | | | | |
| ***T. celericrescens*** | 83.3 | 76.7 | 100.0 | | | | |
| ***T. cleftensis*** | 78.7 | 77.3 | 76.7 | 100.0 | | | |
| ***TkPTP*** | 76.2 | 74.8 | 74.8 | 75.5 | 100.0 | | |
| ***T. gorgonarius*** | 75.3 | 78.7 | 74.7 | 78.7 | 80.3 | 100.0 | |
| ***ShufPTP*** | 82.0 | 86.0 | 82.0 | 83.3 | 84.3 | 90.0 | 100.0 |

**Figure S1. Multiple sequence alignment of protein sequences used for random shuffling generation of ShufPTP**. Sequences were obtained from GenBank.[33] (**A**) comparison between sequences of ShufPTP and its progenitors. Sequences correspond to a dual specificity protein phosphatase from *Thermococcus gorganarius* (WP_088884773.1), *Thermococcus siculi* (WP_088856889.1), *Thermococcus radiotolerans* (WP_088865855.1), *Thermococcus cleftensis* (WP_014789384.1), and *Thermococcus celericrescens* (WP_058937812.1). Residues colored using the Clustal X[34] coloring scheme. Positions that are not conserved across organisms are marked with '*', and also highlighted on (**B**) the structure of ShufPTP. (**C**) Comparison between sequences of ShufPTP and its closest parent, TgPTP, with positions that differ between the two PTPs highlighted on (**D**) the ShufPTP structure. (**Bottom**) The table shows % sequence identity between PTP pairs, calculated using the Standard Protein BLAST (BLASTp) webserver.[35, 36]

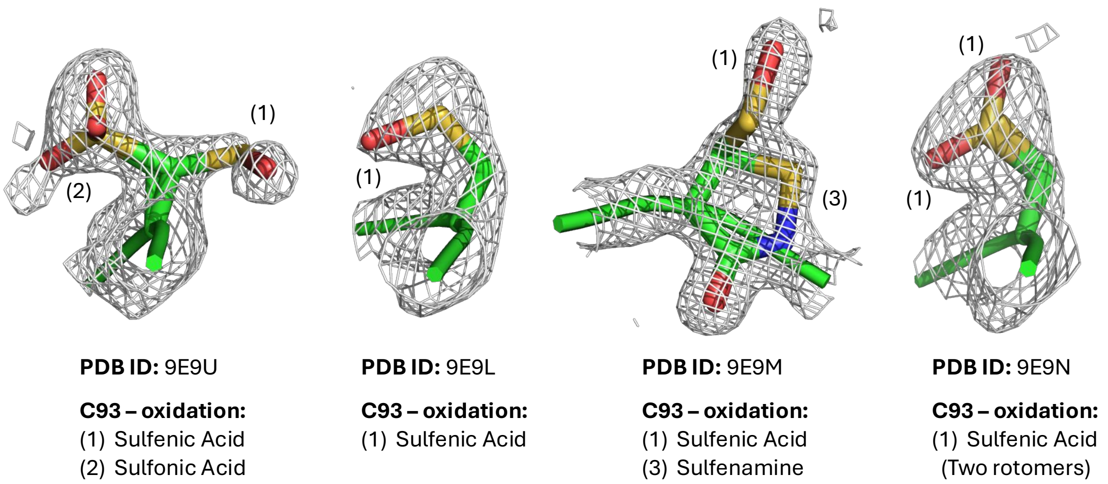


**Figure S2. Simulated annealing omit maps showing oxidized C93 in each of the ShufPTP structures.** Maps are contoured at 1σ. Oxidation states are indicated as described in **Figure 4**.

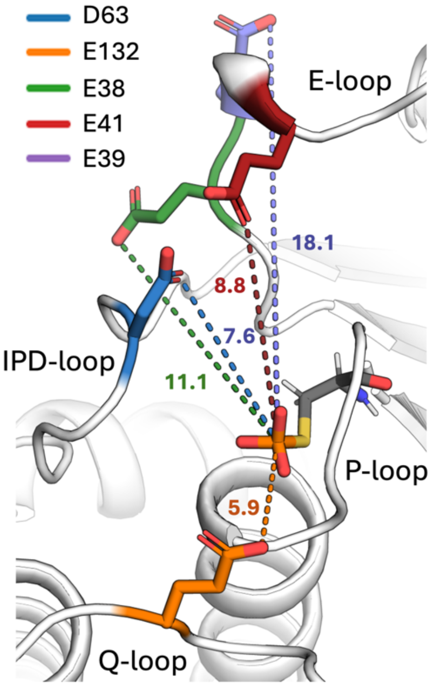


**Figure S3. Prospective catalytic backups in the ShufPTP active site**. Prospective catalytic backup sidechains in the ShufPTP active site were identified based on the distance between the closest oxygen atom of the side chain and the phosphorus atom of the phosphoenzyme intermediate (which was modelled manually into the structure) highlighted, in the crystal structure of wild-type ShufPTP (PDB ID: 9E9N, this work). Note that although the E-loop residues E38 and E39 clearly point away from the active site in the crystal structure of ShufPTP, we nevertheless included these side chains in our analysis, in case rearrangement of the E-loop could bring these residues closer to the active site during our molecular dynamics simulations.

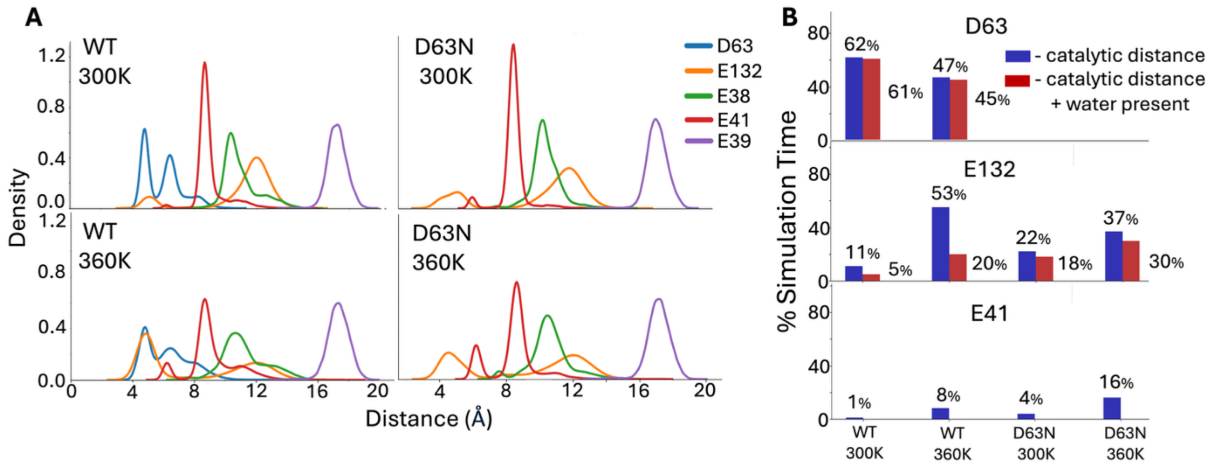


**Figure S4. Identifying prospective catalytic backups in the ShufPTP active site.** Shown here are the distances between the phosphate group and prospective catalytic carboxylate side chains, extracted from our MD simulations of wild-type and D63N ShufPTP, at the respective phosphoenzyme intermediate states. Simulations were performed at both 300 and 360K. (**A**) Kernel density estimates (KDE) of distances between the phosphate group and prospective catalytic side chains, defined as the distance between the P atom of the phosphate group and the closest oxygen atom of the carboxylate side chain of each key residue, were compared to identify a backup catalytic residue. (**B**) The % simulation time the side chains of each of the three most likely candidates (D63, E41 and E132) spent within 6Å of the phosphate group phosphorus atom (shown in blue). Further, to be catalytically viable, it is necessary for a water molecule to bridge the respective side chain and the phosphate group, in order to act as a nucleophile (**Figure 1**). For each prospective catalytic side chain, we calculated the % of simulation time a water molecule is positioned within both 3.5Å of the carboxylic acid of the respective side chain, and of the phosphorus atom of the phosphate group (red, measured based on distances to the nucleophilic oxygen atom). This panel shows that even though there is a small peak at smaller distances for the E41 side chain in panel A, this side chain is essentially never sampling a catalytic conformation on these simulation timescales, and thus the E41 side chain was ruled out from further consideration.

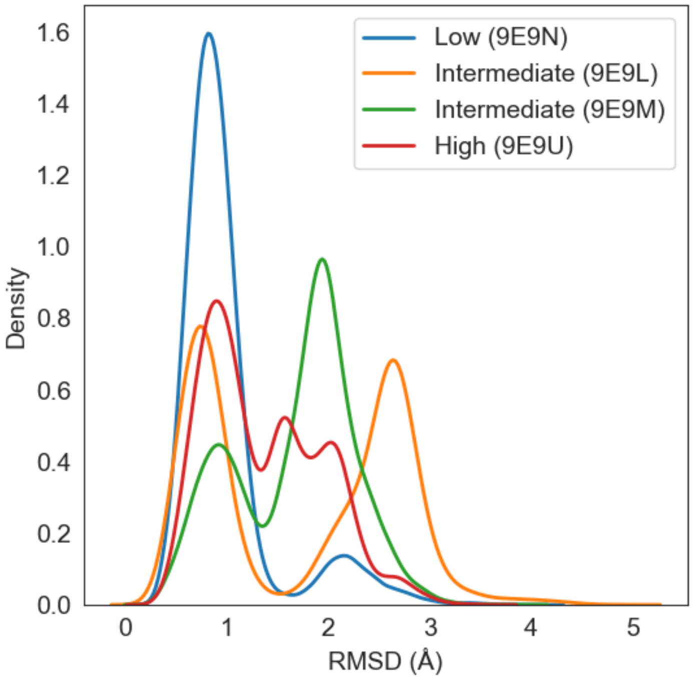


**Figure S5. Geometric stability of the R99 side chain.** The root means square deviation (RMSD, Å) of the R99 side chain in the simulations of unliganded ShufPTP, initialized from all four crystal structures of ShufPTP presented in this work (**Table S1**). The low P-loop crystal structure was chosen as a common reference for all RMSD measurements. While all simulations show some stabilization of the R99, side chain, which is a key residue required for phosphate binding, only simulations of the "low" crystal structure shows the conformation captured in crystal structures as highly preferred.

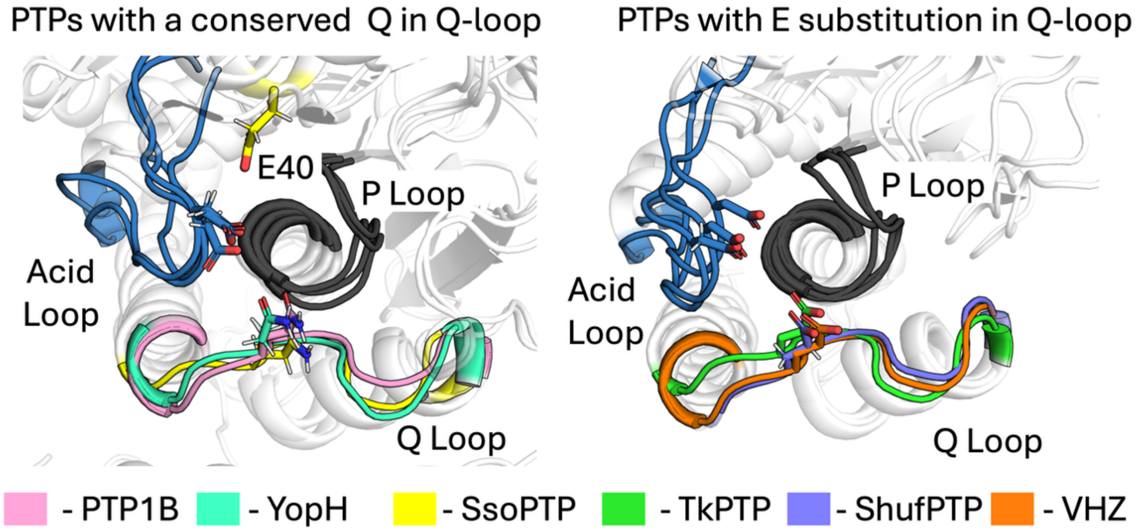


**Figure S6. Q *vs.* E on the Q-loop of PTPs.** Shown here is an overlay of structures of PTPs with (**left**) a conserved Q in their Q-loop (specifically, PTP1B, YopH and SsoPTP), and (**right**) an E in the corresponding position (specifically TkPTP, ShufPTP and VHZ). The latter three enzymes all utilize backup catalytic mechanisms.[2, 37] Shown here is also the position of E40 on the SsoPTP structure, which acts as an alternate catalytic backup in the absence of a Q-loop glutamic acid.[38]

.

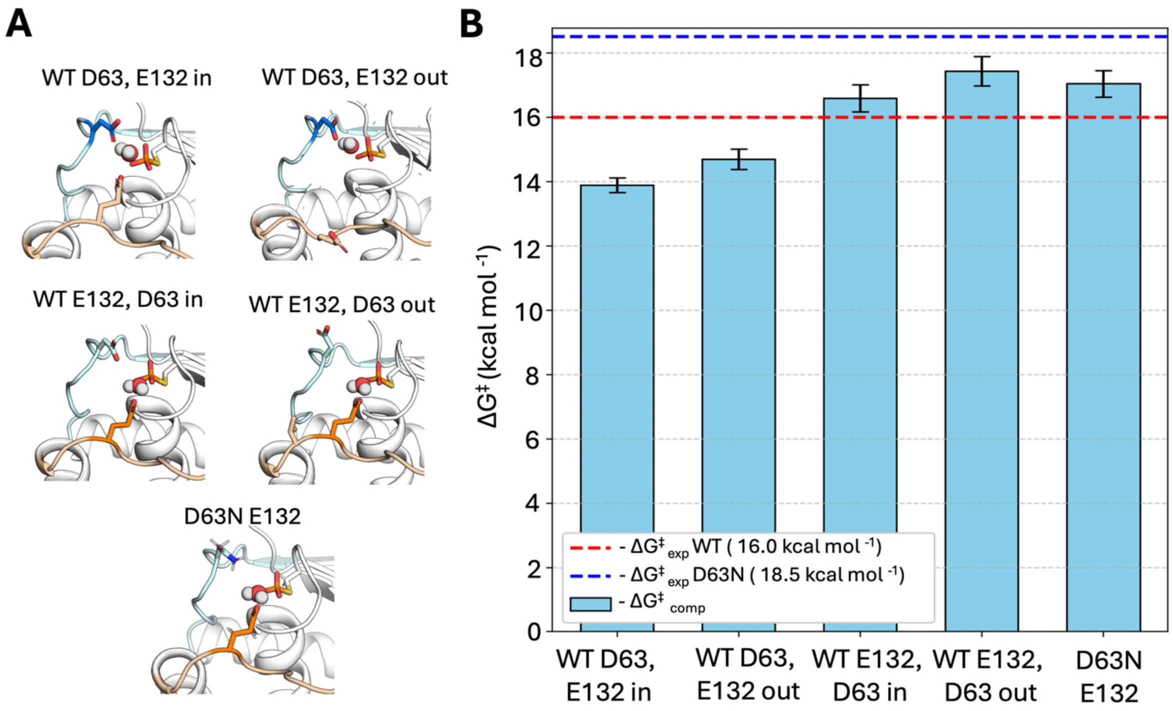


**Figure S7. Active site plasticity and backup catalytic mechanisms in ShufPTP. (A)** Illustration of the reaction environment in five different catalytic scenarios that were simulated using the empirical valence bond (EVB) approach,[23] to evaluate energy barrier of the reaction facilitated by proton transfer to the D63 or E132 side chains in ShufPTP, and the E143 side chain in the D63N variant**.** In these scenarios, either or both of the D63 and E132 side chains can be pointing into ("in") or out ("out") from the ShufPTP active site. **(B)** The calculated $\Delta G^{\ddagger}$ values (kcal $mol^{-1}$) for each scenario were compared to the experimental values obtained from the kinetic data ($k_{cat}$, **Table 2**) for both variants. The red dashed line indicates the experimental activation free energy for the reaction catalyzed by ShufPTP, and the blue dashed line indicates the experimental activation free energy for the reaction catalyzed by the D63N variant. The error bars represent the standard error of the mean on the calculated activation free energies over 20 individual EVB trajectories for each system. The raw data for this figure are shown in **Table S3**.

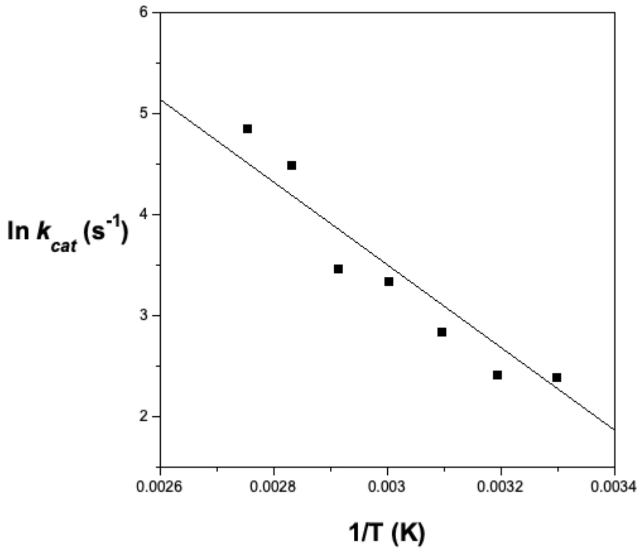


**Figure S8**. **Thermal stability of ShufPTP.** The Arrhenius plot for ShufPTP-catalyzed hydrolysis of *p*NPP in the range of 22 - 90 °C indicates high thermostability up to 90 °C. Loss of activity at high temperatures would result in downward curvature at the highest temperatures (the left side of the graph), which is not observed.

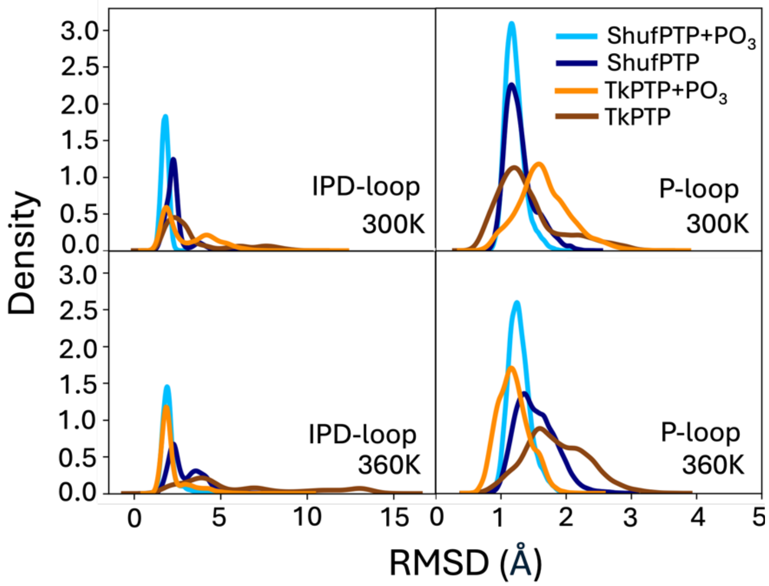


**Figure S9**. **Displacement the IPD- and P-loops of ShufPTP and TkPTP, during MD simulations of active(low) state**. Kernel density estimation (KDE) analysis of the root mean square deviations (RMSD, Å) for the backbone atoms of the P-and IPD-loops in simulations of TkPTP in its active P-loop state and ShufPTP in the low P-loop state. Shown here are data for both unliganded enzymes and the phosphoenzyme intermediate states (+$PO_3$), at 300 and 360K. At the phosphoenzyme intermediate state, the conformational ensembles of the IPD- and P-loops shift closer to the respective P-loop active state crystal structure (PDB IDs: 5Z5A[2] and 9E9N), especially in the case of ShufPTP. When the simulation temperature is increased to 360K a greater flexibility of both loops is observed in the unphosphorylated systems. However, in the case of the phosphorylated system the stability of TkPTP catalytic conformation is increased.

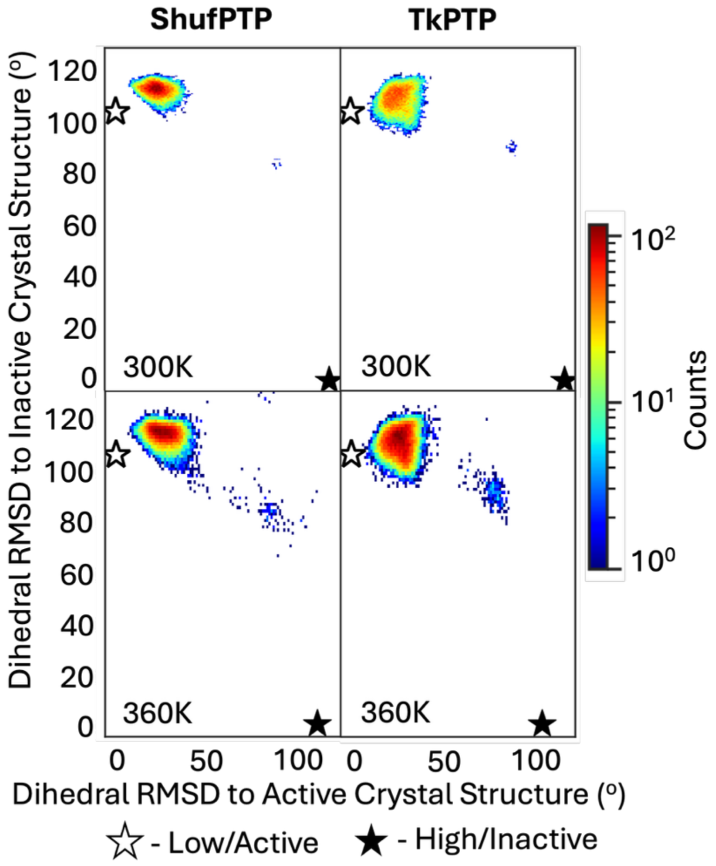


**Figure S10**. **Comparison of P-loop conformational transitions between ShufPTP and TkPTP, in simulations initialized from the active (low) conformation of the P-loop**. Shown here are 2D histograms of the P-loop conformation sampled in our MD simulations of each enzyme. The deviation between active and inactive state X-ray crystal structures (CS) of ShufPTP and TkPTP corresponds to $\varphi$ and $\psi$ angles of 116.1 degrees and 104.7 degrees respectively. Simulations were initiated from the active (low) state of each enzyme, with the IPD-loop in its loop-closed state. The corresponding crystallographic positions of the P-loop are denoted by stars, as annotated on the figure.

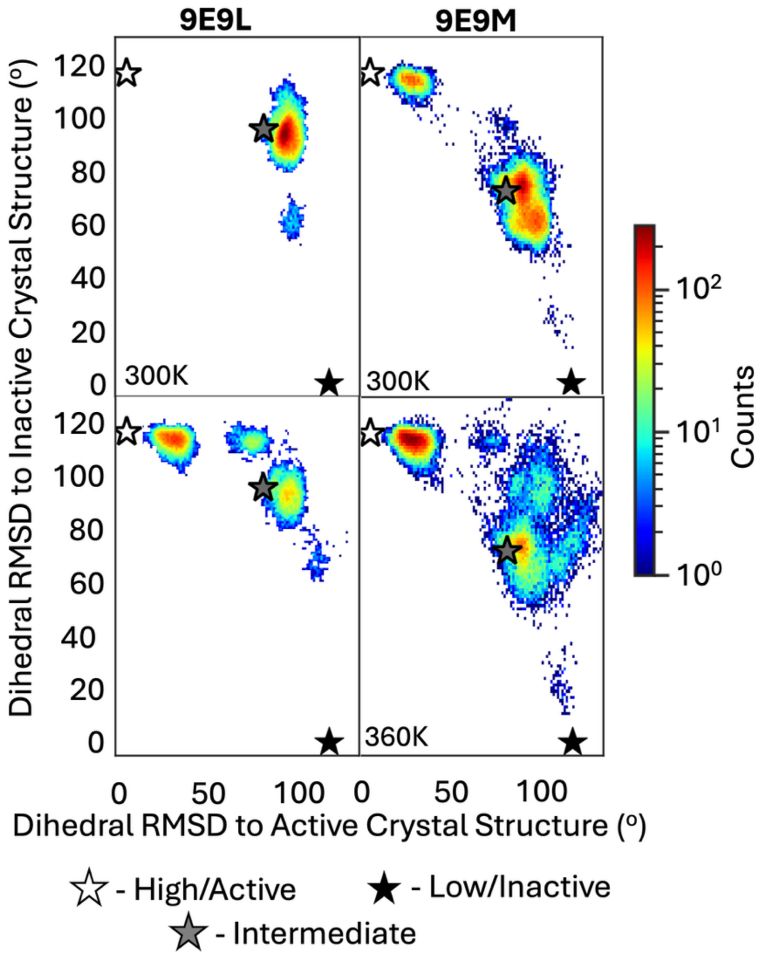


**Figure S11**. **Comparison of P-loop conformational transitions between ShufPTP, in simulations initialized from the intermediate conformations of the P-loop**. Shown here are 2D histograms of the P-loop conformation sampled in our MD simulations of each enzyme. The deviation between low and high state X-ray crystal structures (CS) of ShufPTP correspond to $\varphi$ and $\psi$ angles of 116.1 degrees. The angle RMSD of the two ShufPTP intermediate to active (low) and inactive (high) CS are: 9E9L = (78.6, 95.2) and 9E9M = (78.6, 71.6) degrees. Simulations were initiated from both experimentally obtained crystalized intermediate states of P-loop conformation of ShufPTP (PDB ID: 9E9L and 9E9M), with the IPD-loop in its loop-closed state. The corresponding crystallographic positions of the P-loop are denoted by stars, as annotated on the figure.

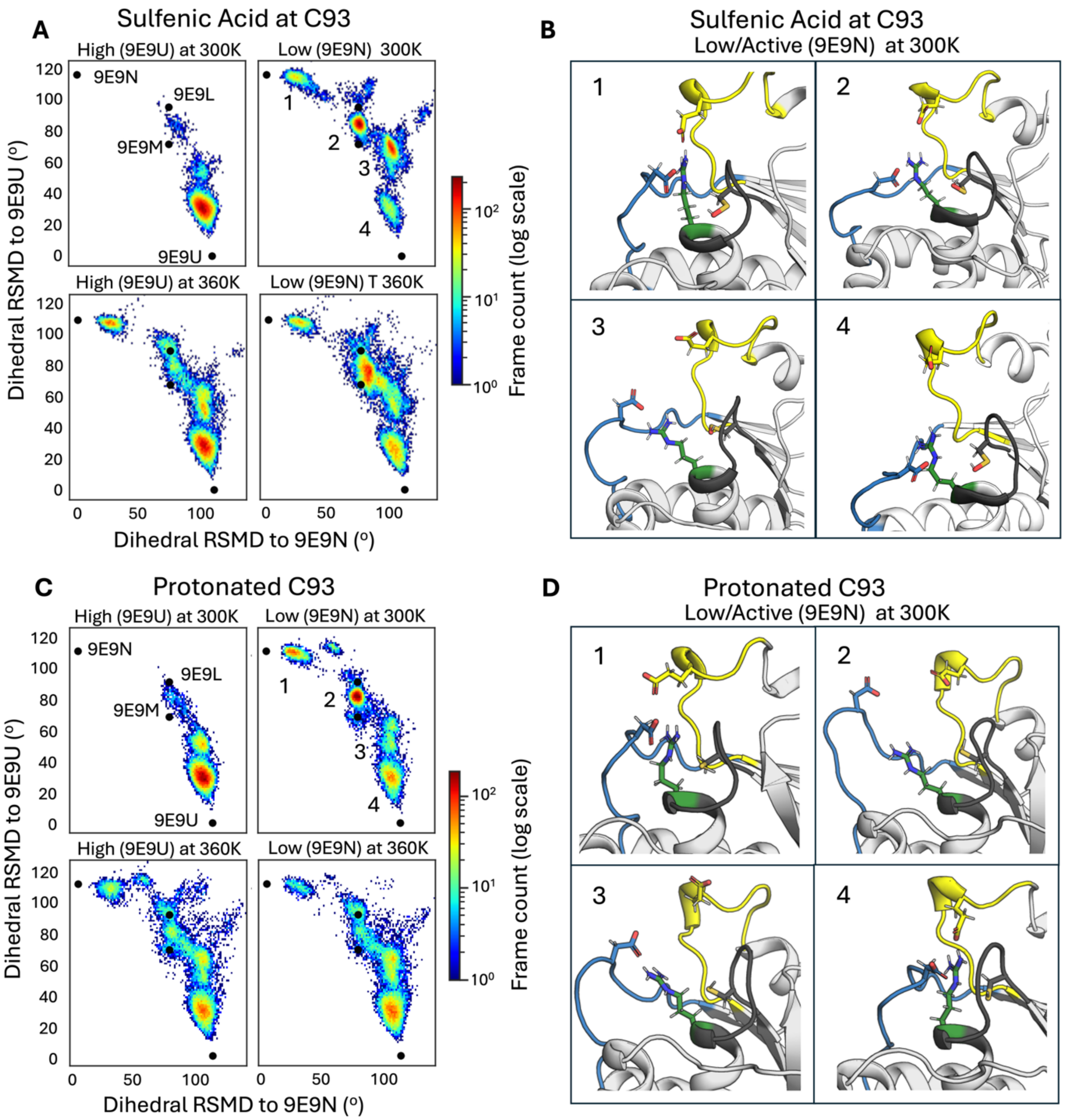


**Figure S12. Comparison of P-loop conformational transitions in ShufPTP with its nucleophilic cysteine side chain in its (A) S-hydroxycysteine and (C) protonated forms.** 2D histograms of the P-loop conformation sampled in MD simulations of ShufPTP where C93 is in its S-hydroxycysteine form. MD simulations were initialized in active/high and inactive/low states P-loop states, and conducted at both 300K and 360K. PDB IDs for the structures used for this analysis are shown on the different panels in parentheses. The dihedral RMSD was defined based on a root mean square deviation (RMSD)-like metric of angles throughout the P-loop (see Eq. 1 of the main text). The deviation between active/low and inactive/high P-loop state X-ray crystal structures of ShufPTP corresponds to dihedral RMSD values of 116.1 degrees and 104.7 degrees respectively. The dihedral RMSD of two ShufPTP intermediates to active/low and inactive/high crystal structure are: 9E9L = (78.6, 95.2) degrees, and 9E9M = (78.6, 71.6) degrees. **(B, D)** Representative snapshots obtained from the low/active P-loop state simulations at 300K in each Cys form were visualized and numbered according to the numbering on the plots A and C. Structures were colored as follows: acid loop in blue, E-loop in yellow and P-loop in black. The catalytically relevant Arg99 residue is shown in green.

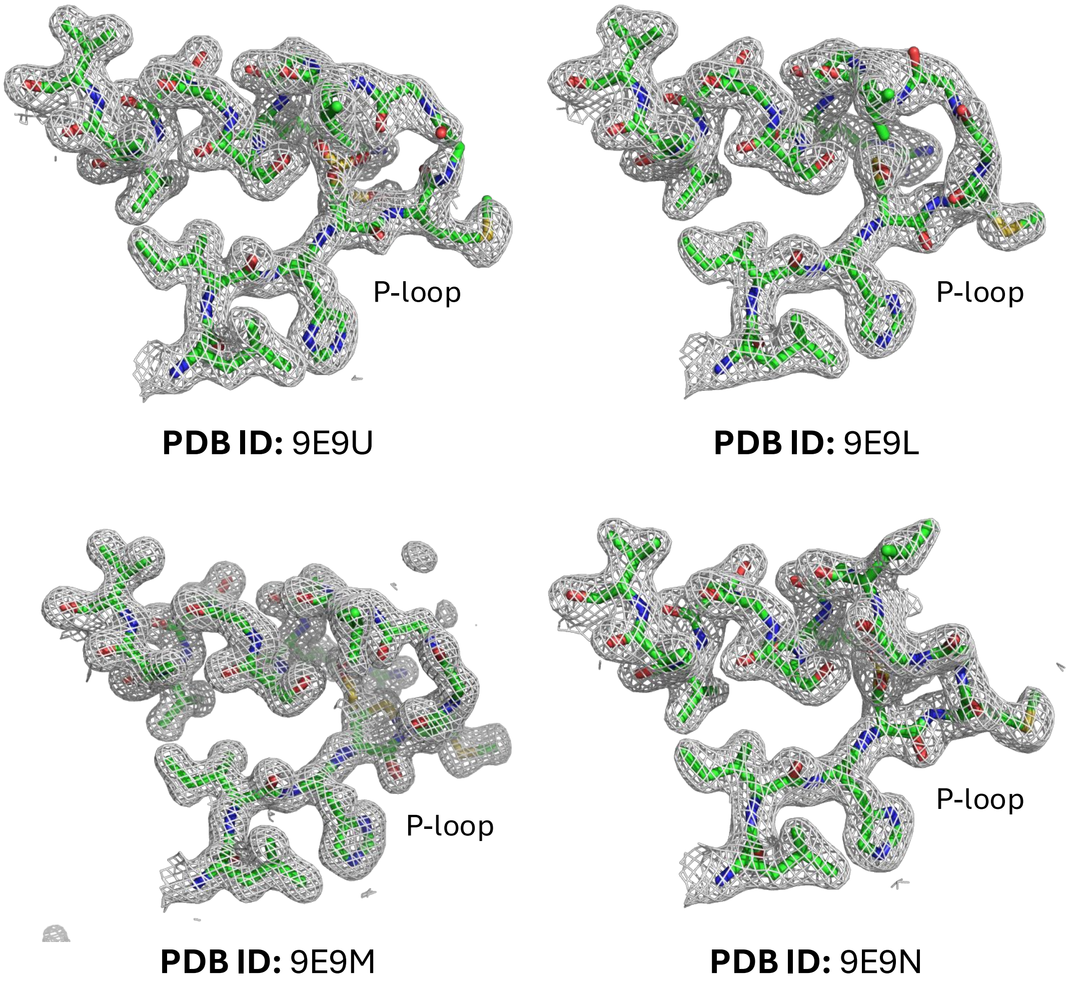


**Figure S13. Representative electron density maps for each of the ShufPTP structures.** 2fo-fc difference maps contoured at 1s. Residues 90-105 (including the P-loop) are shown.

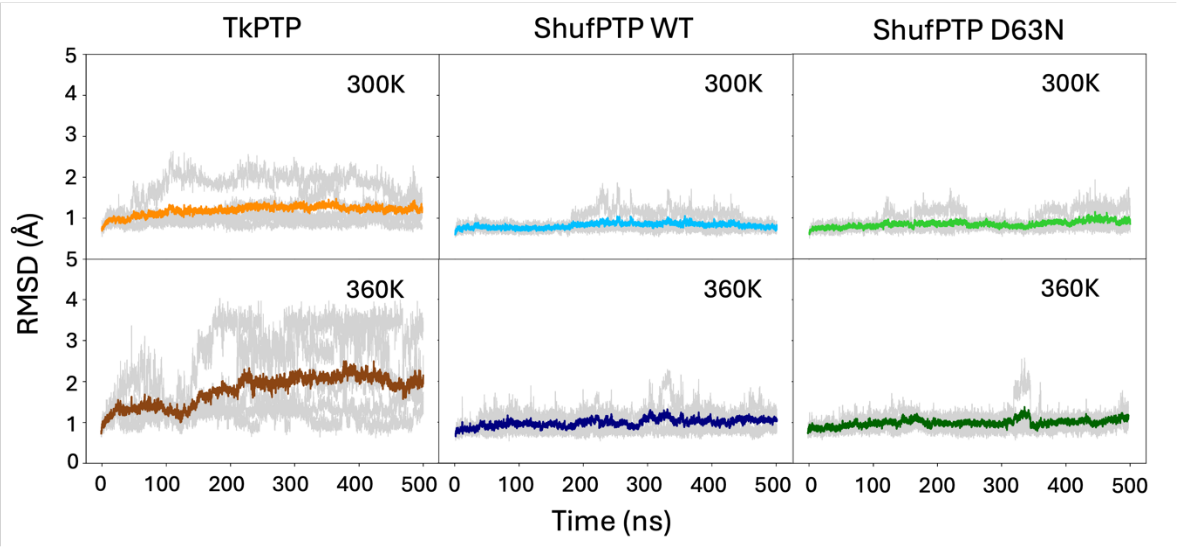


**Figure S14**. Comparison of backbone root mean square deviations (RMSD, Å) of molecular dynamics simulations of wild-type TkPTP, ShufPTP and the ShufPTP D63N variant, initialized from the unliganded active (low) P-loop crystal structure of ShufPTP at 300K and 360K (PDB IDs: 5Z5A,[2] 9E9N).

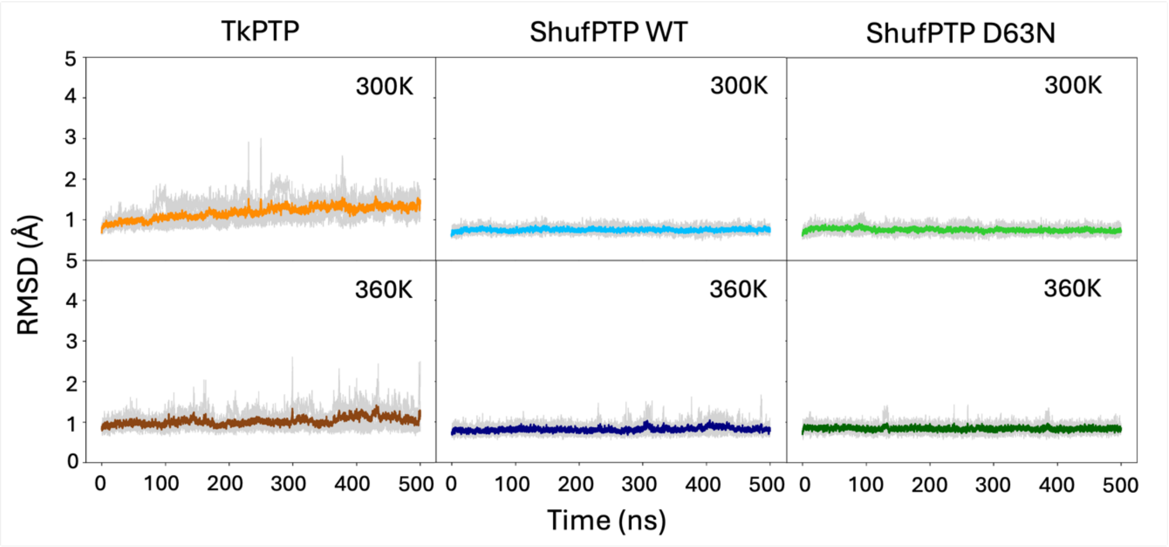


**Figure S15**. Comparison of backbone root mean square deviations (RMSD, Å) of molecular dynamics simulations of wild-type TkPTP, ShufPTP and the ShufPTP D63N variant, initialized from the active (low) P-loop crystal structure of ShufPTP with phosphorylated C93, at 300K and 360K (PDB IDs: 5Z5A,[2] 9E9N).

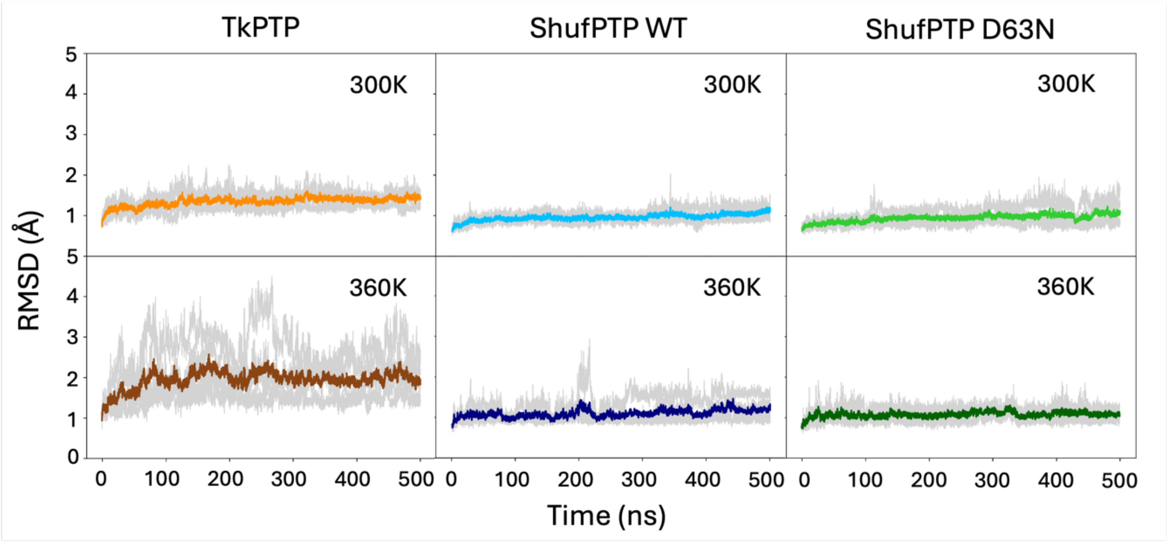


**Figure S16**. Comparison of backbone root mean square deviations (RMSD, Å) of molecular dynamics simulations of wild-type TkPTP, ShufPTP and the ShufPTP D63N variant , initialized from the unliganded inactive (high) P-loop crystal structure of ShufPTP at 300K and 360K (PDB IDs: 5Z59,[2] 9E9U).

### S3. Supplementary Tables

**Table S1.** X-ray data collection and refinement statistics.

| PDB ID | 9E9N | 9E9M | 9E9L | 9E9U |
|---|---|---|---|---|
| P-loop Conformation | Low | Intermediate | Intermediate | High |
| C93 Oxidation State | Sulfenic Acid | Sulfenic Acid<br>Sulfenamide | Sulfenic Acid | Sulfenic Acid<br>Sulfonic Acid |
| Ligand | PEG | Sulfate | Vanadate | Vanadate |
| **Data collection** | | | | |
| Source | Home Source | Home Source | SSRL 9-2 | SSRL 9-2 |
| Wavelength (Å) | 1.5418 | 1.5418 | 0.97946 | 0.97946 |
| Resolution range (Å) | 50.0-1.94<br>(2.01-1.94)* | 50.0-1.55<br>(1.61-1.55) | 50.0-2.10<br>(2.18-2.10) | 50.0-1.80<br>(1.87-1.80) |
| Space group | P $6_1$ | P $6_1$ | P $6_1$ | P $6_1$ |
| Unit cell *a*, *b*, *c* (Å) | 85.64, 85.64, 32.2 | 86.00, 86.00, 32.2 | 85.72, 85.72, 32.37 | 86.05, 86.05, 32.23 |
| $\alpha, \beta, \gamma$ (°) | 90, 90, 120 | 90, 90, 120 | 90, 90, 120 | 90, 90, 120 |
| Total reflections | 182280 | 385923 | 112232 | 213691 |
| Unique reflections | 10023 (860) | 19687 (1907) | 7896 (669) | 12709 (1148) |
| Redundancy | 18.1 (4.9) | 19.6 (16.5) | 14.2 (11.1) | 16.8 (6.6) |
| Completeness (%) | 98.3 (84.7) | 98.2 (95.2) | 96.9 (83.5) | 98.9 (91.3) |
| Mean I/$\sigma$(I) | 26.5 (1.4) | 51.3 (5.9) | 15.53 (5.5) | 17.0 (2.1) |
| Wilson B-factor | 34.6 | 20.8 | 34.1 | 25.5 |
| R merge | 0.100 (0.983) | 0.062 (0.534) | 0.174 (0.451) | 0.168 (0.641) |
| CC1/2 | (0.575) | (0.942) | (0.942) | (0.743) |
| **Refinement** | | | | |
| No. of reflections | 10021 (860) | 19687 (1907) | 7893 (670) | 12709 (1148) |
| $R_{work}/R_{free}$ | 0.178/0.231<br>(0.330/0.471) | 0.185/0.211<br>(0.244/0.334) | 0.184/0.224<br>(0.222/0.234) | 0.178/0.214<br>(0.271/0.328) |
| No. of atoms | | | | |
| Protein | 1199 | 1219 | 1190 | 1221 |
| Ligand/ion | 13 | 10 | 29 | 5 |
| Water | 43 | 129 | 18 | 52 |
| B-factors | | | | |
| Protein | 35.6 | 22.6 | 35.0 | 26.6 |
| Ligand/ion | 45.1 | 27.6 | 53.6 | 39.2 |
| Water | 41.5 | 32.2 | 40.5 | 31.7 |
| R.m.s. deviations | | | | |
| Bond lengths (Å) | 0.011 | 0.011 | 0.011 | 0.011 |
| Bond angles (°) | 1.196 | 1.196 | 1.196 | 1.196 |
| Ramachandran plot | | | | |
| Favored (%) | 89.4 | 93.4 | 89.4 | 89.4 |
| Allowed (%) | 9.5 | 6.1 | 9.5 | 9.5 |
| Outliers (%) | 1.0 | 0.4 | 1.0 | 1.0 |
| Rotamer outliers (%) | 0.82 | 0.00 | 5.65 | 3.88 |
| Clashscore | 4.97 | 7.71 | 7.45 | 2.84 |

*Values in parentheses are for highest-resolution shell.

**Table S2.** Summary of the DelPhiPKa[39] calculated p$K_a$ values of the nucleophilic cysteine across a range of PTPs.[a]

| **System** | $k_{cat}$ **(s$^{-1}$)** | $k_{inact}$ **(M$^{-1}$ s$^{-1}$)** | **Calculated Cysteine p$K_a$** | **Calculated Thiolate Fraction** |
|---|---|---|---|---|
| PTP1B | 52 | 18.5 ± 1.6 | 4.49 ± 0.38 | 0.67 |
| YopH | 720 | 33.0 ± 4.1 | 4.38 ± 0.39 | 0.70 |
| TkPTP (active) | 4.7 | N/A | 5.52 ± 0.26 | 0.14 |
| TkPTP (inactive) | | | 5.60 ± 0.33 | 0.12 |
| ShufPTP (active/low) | 8.6 | 64.8 ± 5.0 | 5.34 ± 0.27 | 0.20 |
| ShufPTP (inactive/high) | | | 5.81 ± 0.38 | 0.08 |

[a] p$K_a$ values were calculated using DelPhiPka, based on 50 snapshots uniformly extracted across 5 x 500 ns MD simulations of each system. In all cases**,** simulations were initialized from structures with closed WPD-loop/IPD-loop conformations. In the case of TkPTP and ShufPTP, multiple P-loop conformations were considered, as shown in parenthesis. Experimental $k_{inact}$ values shown in **Table 1** are presented here for comparison. Experimental $k_{cat}$ values for PTP1B, YopH and TkPTP are presented in **Table 2**. The thiolate fraction was calculated based on the Henderson–Hasselbalch expression, evaluated at a pH of 4.75.

**Table S3.** A comparison of experimental and calculated activation free energies of the hydrolysis of the phosphoenzyme intermediate in the reaction catalyzed by ShufPTP and its D63N variant.[a]

| System | $\Delta G^{\ddagger}_{calc}$ | $\Delta G^{\ddagger}_{exp}$ |
|---|---|---|
| **ShufPTP** | | |
| **D63-as-base, E132 in** | 13.9 ± 0.2 | 16.0 |
| **D63-as-base, E132 out** | 14.7 ± 0.3 | |
| **E132-as-base, D63 in** | 16.6 ± 0.4 | |
| **E132-as-base, D63 out** | 17.4 ± 0.4 | |
| **D63N ShufPTP** | | |
| **E132-as-base** | 17.0 ± 0.4 | 18.5 |

[a] All calculated energies ($\Delta G^{\ddagger}_{calc}$) are presented as average values and standard errors of the mean across 20 independent EVB trajectories, obtained as described in the Materials and Methods. "In" and "out" refer to whether a given side chain is pointing into or out of the active site (**Figure S9**). Note that the catalytic residue is always pointing into the active site. Experimental activation energies ($\Delta G^{\ddagger}_{calc}$) are calculated based on turnover numbers ($k_{cat}$, $s^{-1}$) presented in **Table 2**, using transition state theory. In the case of ShufPTP, the lowest energy pathways for each of the D63-as-base and E132-as-base mechanisms are highlighted in blue.

**Table S4.** List of ionized residues and histidine protonation states in EVB simulations of ShufPTP.[a]

| Type | Residue Number |
|---|---|
| **Asp** | 63 |
| **Glu** | 22, 38, 39, 41, 132, 135, 138 |
| **Arg** | 6, 16, 99, 122, 124, 127 |
| **His-ε** | 58, 92 |

[α] All other ionizable residues not specified in the table were simulated in their neutral state during the simulations, as they fell outside the explicit solvent sphere. His-ε corresponds to a histidine singly protonated on its $N_{\epsilon 2}$ nitrogen atom.

(39) Pahari, S.; Sun, L.; Basu, S.; Alexov, E. DelPhiPKa: Including Salt in the Calculations and Enabling Polar Residue to Titrate. *Prot. Struct. Func. Bioinformat.* **86**, *86*, 1277-1283.
